# Engineering cGAS-agonistic oligonucleotides as therapeutics and vaccine adjuvants for cancer immunotherapy

**DOI:** 10.1101/2023.07.13.548237

**Authors:** Shurong Zhou, Ting Su, Furong Cheng, Janet Cole, Xiang Liu, Bei Zhang, Shaheer Alam, Jinze Liu, Guizhi Zhu

**Affiliations:** Department of Pharmaceutical Sciences, College of Pharmacy; Biointerfaces Institute. University of Michigan. Ann Arbor, MI 48109, USA; Department of Pharmaceutics and Center for Pharmaceutical Engineering and Sciences, School of Pharmacy, Virginia Commonwealth University, Richmond, VA 23298, USA; Department of Biostatistics, School of Medicine; Bioinformatics Shared Resource, Massey Cancer Center; Virginia Commonwealth University, Richmond, VA 23298, USA

**Keywords:** cGAS, oligonucleotide therapeutics, immunostimulant, cancer vaccines, combination immunotherapy.

## Abstract

Current cancer immunotherapy (e.g., immune checkpoint blockade (ICB)) has only benefited a small subset of patients. Cyclic GMP-AMP synthase-stimulator of interferon genes (cGAS-STING) activation holds the potential to improve cancer immunotherapy by eliciting type-I interferon (IFN-I) responses in cancer cells and myeloid cells. Yet, current approaches to this end, mostly by targeting STING, have marginal clinical therapeutic efficacy. Here, we report a cGAS-specific agonistic oligonucleotide, Svg3, as a novel approach to cGAS-STING activation for versatile cancer immunotherapy. Featured with a hairpin structure with consecutive guanosines flanking the stem, Svg3 binds to cGAS and enhances cGAS-Svg3 phase separation to form liquid-like droplets. This results in cGAS activation by Svg3 for robust and dose-dependent IFN-I responses, which outperforms several state-of-the-art STING agonists in murine and human immune cells, and human tumor tissues. Nanocarriers efficiently delivers Svg3 to tissues, cells, and cytosol where cGAS is located. Svg3 reduces tumor immunosuppression and potentiates ICB therapeutic efficacy of multiple syngeneic tumors, in wildtype but neither *cGas*^-/-^ nor goldenticket *Sting*^-/-^ mice. Further, as an immunostimulant adjuvant, Svg3 enhances the immunogenicity of peptide antigens to elicit potent T cell responses for robust ICB combination immunotherapy of tumors. Overall, cGAS-agonistic Svg3 is promising for versatile cancer combination immunotherapy.

## Introduction

ICB immunotherapy has improved the treatment outcomes of a growing number of cancers. However, most of cancer patients have not benefited from current ICB, partly due to immunosuppressive tumor microenvironment (TME), a lack of pre-existing antitumor immune cells, and immune-related adverse events^[1]^. Immunostimulants and cancer antigen-specific vaccines hold a great potential to overcome these challenges and therefore maximize the clinical benefit of ICB for cancer treatment.

As pattern recognition receptors (PRRs), cGAS and STING are emerging targets for the development of immunostimulants in cancer immunotherapy. Indeed, cGAS-STING immunostimulatory pathway is involved in various diseases such as cancer^[2,3]^, autoimmune diseases^[4,5]^, and senescence^[6,7]^. Specifically, cytosolic long double-stranded DNA (dsDNA) activates cytosolic cGAS to synthesize 2’3’-cyclic GMP-AMP (2’3’-cGAMP) from endogenous ATP and GTP. 2’3’-cGAMP binds to and activates STING on endoplasmic reticulum (ER) membrane, resulting in IFN regulatory factor 3 (IRF3) phosphorylation and nuclear factor kappa B (NF-κB) activation, and eventually IFN-I responses^[8]^. IFN-I are critical cytokines for antigen presentation and T cell activation, making cGAS-STING activation appealing to elicit antitumor T cell responses, which are pivotal for cancer immunotherapy^[9,10]^. Specifically, cGAS-STING activation upregulates the expression of proinflammatory chemokines and co-stimulatory molecules, which together promote antigen presentation and T cell priming, resulting in antigen-specific T cell responses; moreover, tumor IFN-I responses promote tumor infiltration of antitumor immune cells^[11]^. This makes it attractive to activate cGAS-STING pathway for cancer combination therapy^[12,13]^.

To this end, STING agonists, ranging from natural or analog cyclic dinucleotides (CDNs) to synthetic compounds, have been tested preclinically and clinically^[14]^. However, current small molecule STING agonists are associated with various limitations. Natural CDNs are susceptible to nuclease degradation (e.g., ectonucleotide pyrophosphatase/phosphodiesterase 1 (ENPP1))^[15]^, which demands complex stability-enhancer modifications of CDNs to enhance their biostability. Further, CDNs are very hydrophilic and small (∼700 Da) with negative electrostatic charge. This makes it difficult for current drug delivery technologies to efficiently deliver CDNs into target tissues, cells, and cytosol, where STING resides^[16]^. As a result, the antitumor responses and tumor therapeutic efficacy of CDNs, such as a modified CDN ADU-S100 tested in a Phase I clinical trial, has been marginal^[17]^. Non-CDN small molecules are another class of STING agonists under development. However, DMXAA, a preclinical STING agonist, failed to benefit cancer therapy in a Phase III clinical trial due to its selective activation of murine STING, but not human STING^[18]^. More recent non-CDN STING agonists, such as diABZI, showed promising preclinical tumor therapeutic efficacy^[19]^, though their clinical efficacy and safety remain to be evaluated. Finally, STING agonists can be subject to the restriction of human alleles^[16]^. Overall, these challenges call for innovative approaches to drug development for cGAS-STING activation.

To this end, here, we report the engineering of oligonucleotide agonists for cGAS as a novel approach to cGAS-STING immunostimulation for combination cancer immunotherapy. cGAS can be activated by natural dsDNA in a sequence-independent and length-dependent manner. Specifically, cGAS activation requires relatively long dsDNA with length > 45 base pairs (bp) to form ladder-like enzymatically active cGAS dimers^[20,21]^. Interestingly, shorter dsDNA with guanosine (G)-rich overhangs also activate cGAS ^[22]^. Inspired by this, here, by structural engineering and screening a series of hairpin-shaped single stranded DNA (ssDNA), we identified Svg3 as a potent cGAS agonist for versatile applications in combination cancer immunotherapy (**Fig. 1**). Svg3 has a core structure of a hairpin with a GGG triplet in each of the four overhangs adjacent to the hairpin stem. Svg3 was easily synthesized on automated DNA synthesizers, and was efficiently formulated in well-established nucleic acid nanocarriers, such as liposomes and lipid nanoparticles (LNPs), which allow efficient tissue, cell, and cytosolic delivery of Svg3. Svg3 elicited cGAS-dependent and cGAS-selective IFN-I responses, with undetectable inflammasome activation and pyroptosis, which could also be activated by long dsDNA. Svg3 outperformed several current state-of-the-art STING agonists to induce potent IFN-I responses in a dose-dependent manner in both murine and human immune cells and cancer cells, as well as human tumor tissues. Intratumoral (i.t.) administration of Svg3-loaded nanoparticles significantly reduced TME immunosuppression. In multiple poorly immunogenic syngeneic murine tumor models, i.t. administration of Svg3 nanoparticles significantly enhanced the therapeutic efficacies of ICB. Impressively, Svg3 outperformed several state-of-the-art STING agonists for tumor therapy. Such tumor therapeutic efficacy is cGAS- and STING-dependent, as verified in cGAS- or STING-knockout mice. Moreover, in antigen presenting cells (APCs), as a vaccine adjuvant, Svg3 enhanced the expression of costimulatory factors and the production of proinflammatory cytokines, both of which, along with antigenic epitopes presented on major histocompatibility complexes (MHC), are essential for APCs to license antigen-specific T cells. In mice, subcutaneous (s.c.) administration of Svg3/antigen-loaded nanoparticles promoted their accumulation in draining lymph nodes, resulting in potent antigen-specific T cell responses. As a result, nanoparticulate co-delivery of Svg3 and tumor-specific antigens enhanced ICB therapeutic efficacy of tumor in syngeneic mice. Collectively, these results demonstrated the great potential of Svg3 as a novel, potent, and versatile immunostimulant for combination cancer immunotherapy.

**Figure 1.**
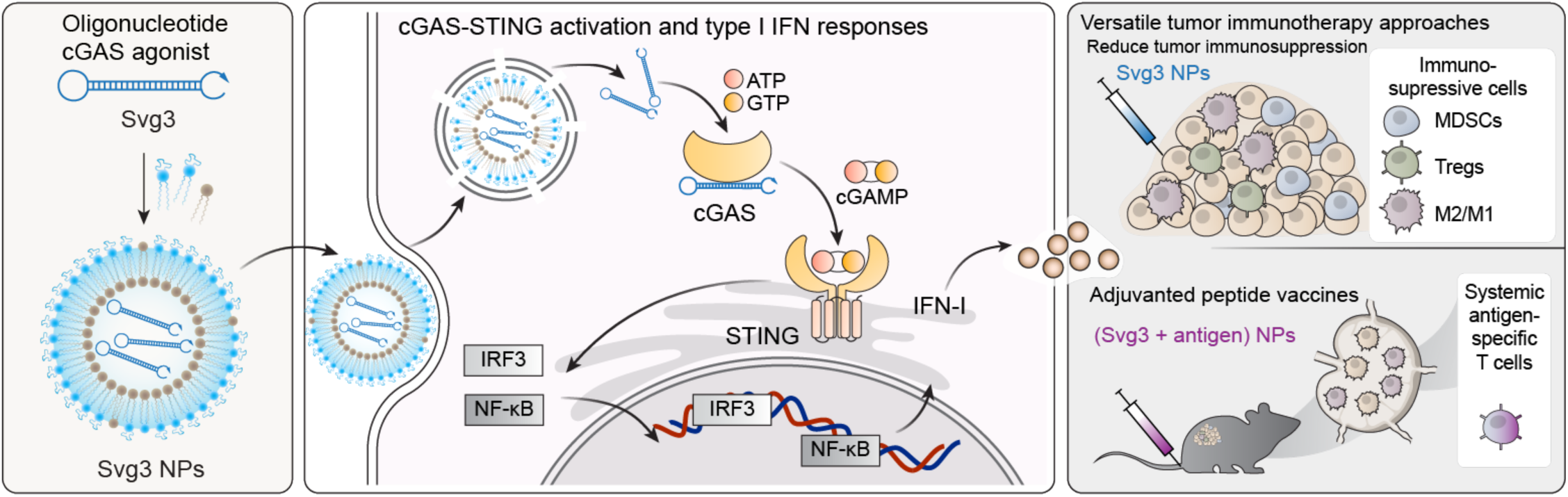
Schematic illustration of an oligonucleotide-based cGAS agonist for cancer immunotherapy. A hairpin-shaped oligonucleotide, named Svg3, is engineered to selectively activate cGAS, and thereby eliciting type I IFN responses in mouse and human cells. Svg3 is readily loaded into well-established nanocarriers for efficient delivery into cells and, upon endosome escape, into cytosol where Svg3 binds to cytosolic cGAS. This results in cGAS-Svg3 phase separation and the formation of cGAS-Svg3 liquid-like droplets, in which cGAS is activated by Svg3 to trigger the activation of cGAS-STING signaling pathway and IFN-I responses. In tumor microenvironment, Svg3-loaded nanoparticles (NPs) reduced tumor immunosuppression and enhanced antitumor immunity. Intratumoral vaccination of Svg3 NPs dramatically potentiated the tumor immunotherapeutic efficacy of ICB in multiple syngeneic murine tumor models. Moreover, as an immunostimulant adjuvant co-loaded with peptide antigens in NPs (Svg3 + antigen) NPs, Svg3 potentiated the immunogenicity of peptide antigens to elicit antigen-specific T cell responses for potent tumor combination immunotherapy.

## Results and discussion

### Identification of cGAS-agonistic Svg3 by oligonucleotide engineering and screening

Inspired by the ability of natural dsDNA to activate cGAS, we attempted to develop oligonucleotide therapeutics as cGAS agonists for application in combination cancer immunotherapy. Relative to dsDNA, ssDNA is expected to benefit from the simplicity of pharmaceutical manufacturing and quality control. Thus, to select potent cGAS-activating ssDNA, we designed a series of hairpin-structured ssDNA oligonucleotides (**Fig. 2A**) by engineering the hairpin stem length, overhang length, as well as consecutive G numbers and positions in overhangs. These ssDNAs were transfected into RAW 264.7 murine macrophages for 24 h, followed by measuring IFN-I response. As a result, oligonucleotide with dsDNA stem length < 16 bp was insufficient to triggering IFN-I response unless consecutive (G)s were added in the overhangs (**Supplementary Fig. 1A**). Elongating the dsDNA stem from 10 bp to 24 bp, which was expected to enhance their cGAS binding affinity, indeed promoted their IFN-I responses, which plateaued at a stem length of 21 bp (**Fig. 2B**). Further elongating the loop or adding G in the overhangs had minimal effect on IFN-I responses (**Fig. 2C, Supplementary Fig. 1B**). Svg3 showed a strong binding affinity with cGAS, with a *Kd* value of 262 ± 14 nM as measured by by microscale thermophoresis (MST) (**Fig, 2D**). As a result, mixing Svg3 with human cGAS protein resulted in the phase separation of cGAS-Svg3 and the formation of cGAS-Svg3 liquid-like droplets (**Fig. 2E**). Moreover, in mouse bone marrow-derived macrophages (BMDMs) and bone marrow-derived dendritic cells (BMDCs), as low as 25 nM Svg3 elicited significant IFN-β production (treatment: 24 h) (**Fig. 2F**). Taken together, Svg3 was the oligonucleotide that elicited the strongest IFN-I responses, and was therefore selected for further studies. Svg3 was predicted to form a hairpin secondary structure with a 21-bp dsDNA stem, a 9-nucleotide loop, and a GGG triplet adjacent to the stem in each of its four overhangs. Mutating the consecutive G to cytosines (C) in Svg3 overhangs dramatically reduced the IFN-I response (**Supplementary Fig. 1C**), validating their critical roles for cGAS activation. Svg3 elicited comparably potent IFN-I responses relative to interferon stimulatory DNA (ISD), a benchmark cGAS-activating 45-bp dsDNA (**Fig. 2G**). We further validated that Svg3 treatment in RAW 264.7 macrophages resulted in efficient 2’3-cGAMP production (**Fig. 2H**) for at least 8 h, which outperformed ISD control whose 2’3’-cGAMP production dramatically reduced to basal level 6 h after treatment. The efficient 2’3’-cGAMP production enabled by Svg3 is consistent with its ability to elicit potent IFN-I responses.

**Figure 2.**
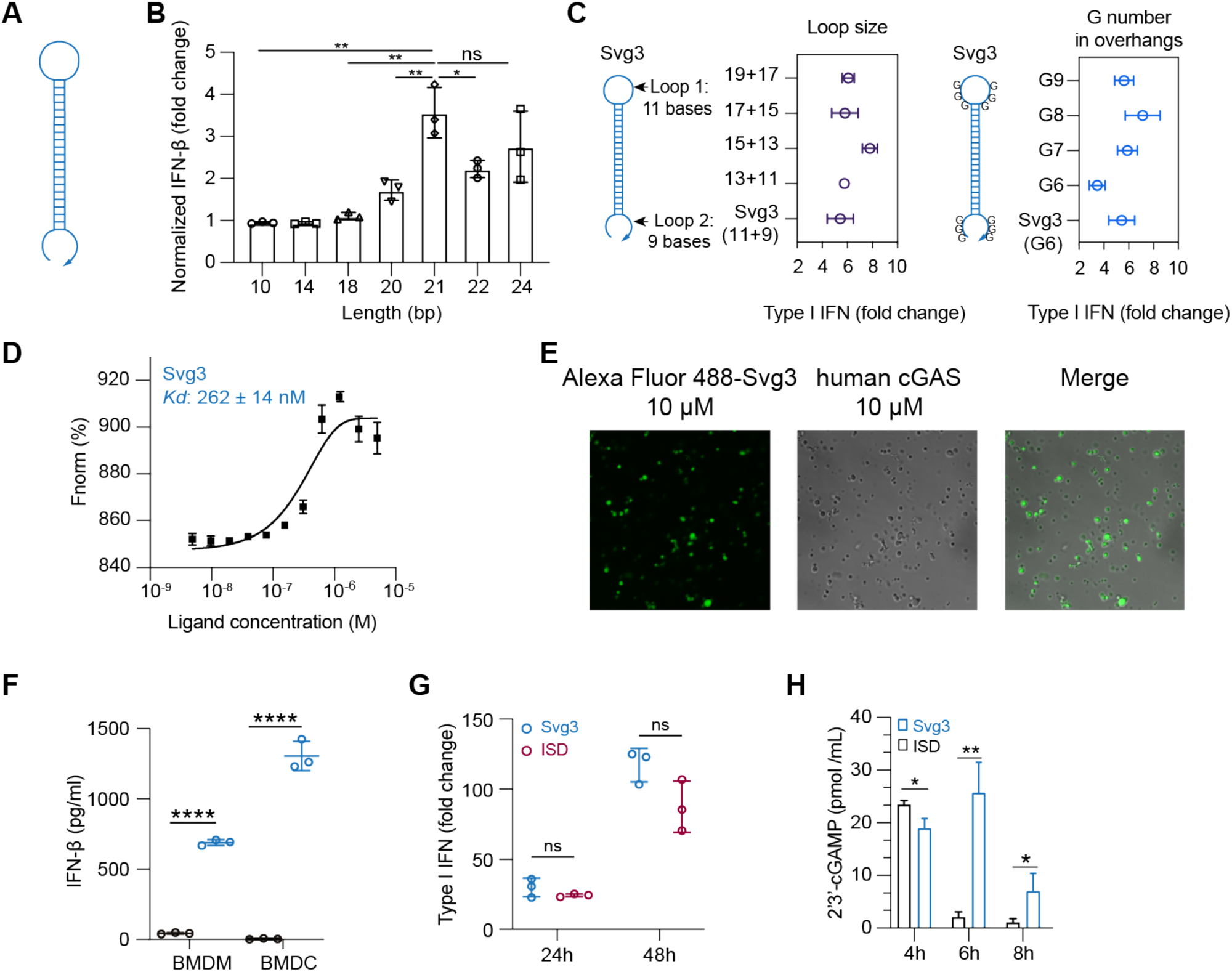
Identification of Svg3 as a potent cGAS agonist by oligonucleotide engineering and screening. **A)** Schematic structures of ssDNA oligonucleotides. **B)** Elongating dsDNA stem length from 10 bp to 24 bp elevated IFN-I response in RAW-ISG macrophages, which plateaued at 21 bp in the stem. **C)** Optimization of the loop sizes and G numbers in hairpin overhangs for cGAS-mediated IFN-I responses in RAW-ISG macrophages. **D)** Svg3 showed a strong binding affinity with human cGAS as measured by MST. **E)** The binding of cGAS with Svg3 induced cGAS-Svg3 phase separation to form liquid-like droplets. Alexa Fluor 488-labeled Svg3 was mixed with human cGAS for 30 min. **F)** Svg3 elicited strong IFN-I responses in murine BMDMs and BMDCs. **G)** Svg3 elicited comparably potent IFN-I responses relative to ISD in RAW-ISG macrophages. **H)** 2’3‘-cGAMP production by RAW 264.7 cells upon treatment with Svg3 or ISD (100 nM) as a control for 4-8 h. 2’3’-cGAMP concentration in cell lysis was measured by ELISA. Unless denoted otherwise, 25 nM DNA was transfected into cells by lipofectamine 2000, followed by incubation for 24 h in cell culture medium before IFN-I measurement. Data: mean ± S.D. p values were determined by t-test (ns: not significant, *p* > 0.05; \**p* < 0.05, \*\**p* < 0.01; \*\*\**p* < 0.001; \*\*\*\**p* < 0.0001).

Oligonucleotide therapeutics are subject to nuclease degradation that limits their biostability and therapeutic efficacy. We verified that Svg3 has decent biostability as shown by the integrity of Svg3 after incubation in PBS and cell culture medium containing 10% fetal bovine serum (FBS) (37°C, 24 h), respectively (**Supplementary Fig. 1D**). We attempted to further prolong the biostability of Svg3 by three approaches. First, we ligated the two terminals of Svg3 ssDNA into circular Svg3 that was expected to avoid exonuclease degradation, however this did not significantly improve its ability to elicit IFN-I in RAW 264.7 macrophage (**Supplementary Fig. 1E**). Second, a phosphonothioate (PS) backbone, which is widely used in oligonucleotide therapeutics to resist nuclease degradation, in the dsDNA stem of Svg3 restricted Svg3’s ability to elicit IFN-I response in RAW 264.7 macrophages (**Supplementary Fig. 1F**). Third, adding nuclease-resistant inverted dT on the 3’ end of Svg3 did not improve the long-term IFN-I response in RAW 264.7 macrophages either (**Supplementary Fig. 1F**). Overall, though further comprehensive testing of modifications may further enhance Svg3 biostability, unmodified Svg3 has already shown great biostability and potent cGAS activation, and was used in the following studies.

To elicit IFN-I responses for cancer immunotherapy *in vivo*, Svg3 is expected to be delivered to immune cells or tumor cells in tumor or lymphoid tissues (e.g., lymph nodes). We leveraged pre-existing drug delivery systems, such as liposomes and LNPs, that have been established for oligonucleotide therapeutics and vaccines. Svg3 was efficiently loaded into liposomes or LNPs (**Supplementary Fig. 2A&B**). Nanoparticles dramatically enhanced the cell uptake of Cy3-labeled Svg3 in RAW 264.7 macrophages (**Supplementary Fig. 2C&D**), followed by endosome escape of Svg3 to reach cytosol, which allows Svg3 to activate cytosolic cGAS (**Supplementary Fig. 2E**). Moreover, liposomes prolonged the tumor retention of i.t. injected Svg3 in 4T1 mammary carcinoma in syngeneic Balb/c mice for at least 7 days (**Supplementary Fig. 2F&G**). These results demonstrated the ability of liposomes to prolong the tissue retention and facilitate cell uptake and cytosolic delivery of Svg3.

### Svg3 is a cGAS-specific agonist

A variety of cytosolic nucleic acid sensors can be activated to elicit IFN-I responses. Specifically, aside from cGAS, cytosolic DNA sensors such as absent in melanoma 2 (AIM2), DEAD box helicase 41 (DDX41), interferon gamma inducible protein (IFI16), Z-DNA binding protein (ZBP1), RNA polymerase III, and LRR binding FLII interacting protein 1 (LRRFIP1) also elicit IFN-I response^[23]^. Moreover, DDX41, IFI16, and ZBP1 induce IFN-I responses in a cGAS-independent and STING-dependent manner^[24]^. To study the cGAS specificity of Svg3-induced IFN-I responses, first, by RNA sequencing (RNA-seq), we analyzed the transcriptomic changes in RAW 264.7 macrophages transfected with Svg3, with blank lipofectamine 2000 as a control. RNA-seq results verified that Svg3 induced a series of ISG, including *Ifna2*, *Ifnb1*, and *Ifit1* (**Fig. 3A**). Interestingly, Svg3 did not significantly upregulate the expression of *Il-1β* (encoding interleukin-1β (IL-1β)) and *Il-18* (**Fig. 3A**). This rules out AIM2 activation by Svg3, which would otherwise trigger confounding inflammasome responses, including IL-1β and IL-18 production. This is further supported by the undetectable IL-1β and IL-18 production by Svg3 in RAW 264.7 macrophages (**Fig. 3B**). Next, using RAW 264.7 macrophages with cGAS knockout (RAW-Lucia™ ISG-KO-cGAS), we then verified that the Svg3-elicited IFN-I responses were cGAS-dependent (**Fig. 3C**). Taken together, these results demonstrated that Svg3 elicited cGAS-specific IFN-I responses.

**Figure 3.**
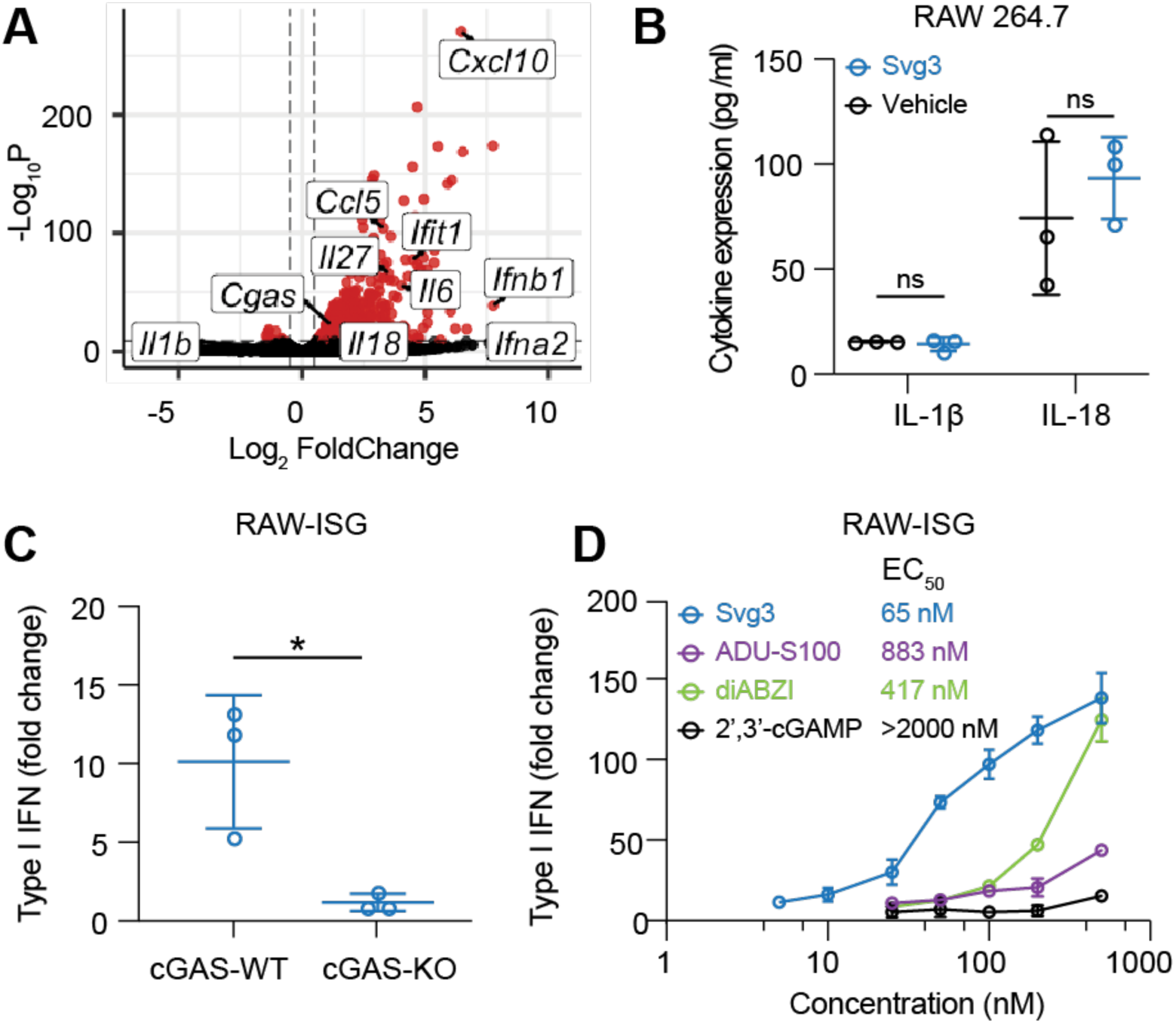
Svg3 specifically activates cGAS to induce potent IFN-I response. **A**) Volcano plot of gene expression in RAW 264.7 macrophages treated with liposomal Svg3 (100 nM Svg3, 6 h). Blank liposome was used as a control. **B**) IL-1β and IL-18 productions in RAW 264.7 macrophages treated with Svg3 (100 nM, 24 h). **C**) IFN-I responses elicited by Svg3 (25 nM, 24 h) in RAW-ISG cells with wildtype cGAS (cGAS-WT) and cGAS knockout (cGAS-KO). **D**) Svg3 was benchmarked against several state-of-the-art STING agonists to elicit IFN-I responses in murine RAW-ISG macrophages. Svg3 outperformed these STING agonists to elicit IFN-I responses. Marked on the right of legends are EC50, indicating that Svg3 showed 13.6x, 6.4x, and 43x lower EC50 than ADU-S100, diABZI, and 2’3’-cGAMP, respectively in RAW-ISG cells. Data: mean ± S.D. *p* values were determined by t-test (ns: not significant, *p* > 0.05; \**p* < 0.05.).

Next, by comparing the IFN-I responses in RAW 264.7 macrophages, we benchmarked Svg3 against multiple types of state-of-the-art STING agonists, including two CDNs 2’3’-cGAMP and ADU-S100, the latter of which is a derivative of c-di-AMP with a strong STING binding affinity, as well as a next-generation non-CDN small molecule diABZI. As a result, Svg3 elicited dose-dependent IFN-I responses with a half maximal effective concentration (EC_50_) as low as 65 nM that outperformed all these STING agonists (**Fig. 3D**). This validated the great potency of cGAS agonist Svg3, making it promising to serve as potent immunostimulant therapeutics and vaccine adjuvants. Overall, these data suggest the ability of Svg3 to elicit potent IFN-I responses.

### Svg3 elicited potent IFN-I responses in human cells and human tumor tissues

We envision that local activation of intratumoral cGAS would promote multifaceted antitumor innate and adaptive immunity. To this end, clinical gene transcript analysis showed high transcript levels of cGAS and STING, respectively, in many types of human tumors such as breast cancer, melanoma, and head and neck cancer (**Supplementary Fig. 3**). This suggests the clinical potential of Svg3 to activate cGAS-STING in the corresponding types of human tumors. The inter-species differences in the immune system represents a barrier for the clinical translation of preclinically tested immunotherapeutics and vaccines. For example, due to slight structural differences of human and murine STINGs, a preclinically promising murine STING agonist, DMXAA, failed in a Phase III clinical trial^[18]^. Further, there are structural differences between mouse cGAS (m-cGAS) and human cGAS(h-cGAS). Relative to m-cGAS, h-cGAS shows reduced binding ability for short dsDNAs (<45 bp)^[21]^. Therefore, it is critical to validate the ability of cGAS agonists to activate h-cGAS and elicit IFN-I responses in human systems. To this end, we first verified that Svg3 elicited potent IFN-I responses, with an EC50 as low as 15 nM in THP-1 human monocytes (**Fig. 4A**). By benchmarking Svg3 against STING agonist 2’3’-cGAMP, Svg3 outperformed 2’3’-cGAMP to elicit IFN-I responses in THP-1 human monocytes (**Fig. 4A**). Lastly, we tested the ability of liposomal Svg3 to elicit IFN-I responses in cultured surgically collected human head and neck squamous cell carcinoma (HNSCC) tissues. We treated these tissues with liposomal Svg3 (0.1 nmole Svg3) for 24 h, followed by quantifying the IFN-I response-associated gene transcripts by quantitative PCR (qPCR). As a result, Svg3 significantly upregulated IFN-I genes in these tissues (**Fig. 4B**). These results highlight the potential of Svg3 for clinical translation in human patients.

**Figure 4.**
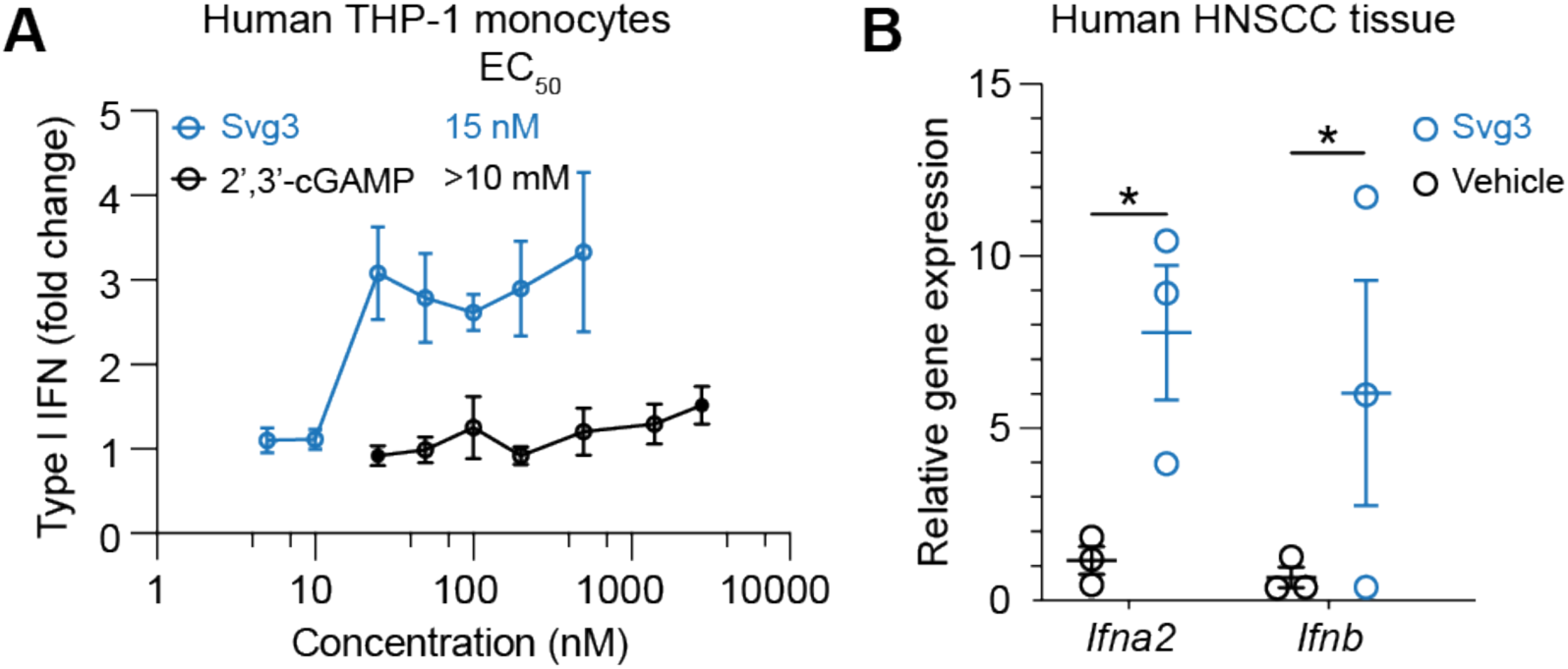
Svg3 elicited potent IFN-I responses in human monocytes and ex vivo human tumor tissues. **A)** Svg3 elicited potent IFN-I responses that outperformed STING agonist 2’3’-cGAMP in human THP-I monocytes. Marked on the right of treatment legends are EC50. Svg3 and 2’3’-cGAMP were transfected using lipofectamine 2000. **B)** Relative gene expression showed that Svg3 elicited IFN-I responses in *ex vivo* HNSCC tissues. Liposomal Svg3 (Svg3: 100 nM) were injected directly into cultured tumor tissues (cohort study, *n* = 3) cultured in medium (1 mL). 24 h later, the mRNA levels of related cytokines were determined by qPCR. Data: mean ± S.D. *P* values were determined by t-test (ns: not significant, *p* > 0.05; \**p* < 0.05.).

### Svg3 reduced TME immunosuppression

TME is primarily where tumor cells suppress antitumor immunity. TME immunosuppression inhibits the tumor infiltration of antitumor immune cells and suppresses the antitumor immunity of intratumoral immune cells. Overcoming TME immunosuppression represents a hallmark to tumor immunotherapy, including the combination of immunostimulants with ICB such as αPD-1. Therefore, we studied TME immunomodulation by Svg3 that elicited potent antitumor IFN-I responses, alone or combined with αPD-1 (i.p. injection) that can reinvigorate antitumor immune cells exhausted by prolonged exposure of inflammatory responses. Motivated by the long tumor retention of liposomal Svg3, we used liposomal Svg3 using 4T1 mammary carcinoma as a model in syngeneic Balb/c mice. Tumor (ca. 50 mm^3^)-bearing mice were treated with liposomal Svg3 (i.t. injection) and/or αPD-1 (i.p. injection), every three days for three times. Three days after the last treatment, tumors were excised to analyze the immune cell composition among all CD45^+^ leukocytes by flow cytometry. Myeloid derived suppressor cells (MDSCs) and regulatory T cells (Treg) are two of the most dominant immunosuppressive cell subsets in TME. Liposomal Svg3, especially when combined with αPD-1, significantly reduced the frequencies of MDSCs and CD4^+^Foxp3^+^CD25^+^ Treg (**Fig. 5B&C**). Meanwhile, liposomal Svg3, alone or combined with αPD-1, promoted the tumor infiltration of CD8^+^ T cells, and enhanced the ratio of CD8^+^ T cells over CD4^+^Foxp3^+^CD25^+^ T cells (**Fig. 5D-F**), both of which are expected to promote tumor therapeutic efficacy and predict tumor therapeutic efficacy. Moreover, liposomal Svg3 + αPD-1 enhanced the frequency of natural killer (NK) cells in TME and promoted the polarization of regulatory M2-like macrophage to proinflammatory M1-like macrophages, as measured M1/M2 ratio (**Fig. 5G&H**). None of these treatments significantly impacted the densities of TME DCs (**Supplementary Fig. 4**). Taken together, liposomal Svg3, especially when combined with αPD-1, reduced multi-tier immunosuppressive cell densities and enhanced the antitumor immune cell densities in TME, which are expected to benefit tumor immunotherapy.

**Figure 5.**
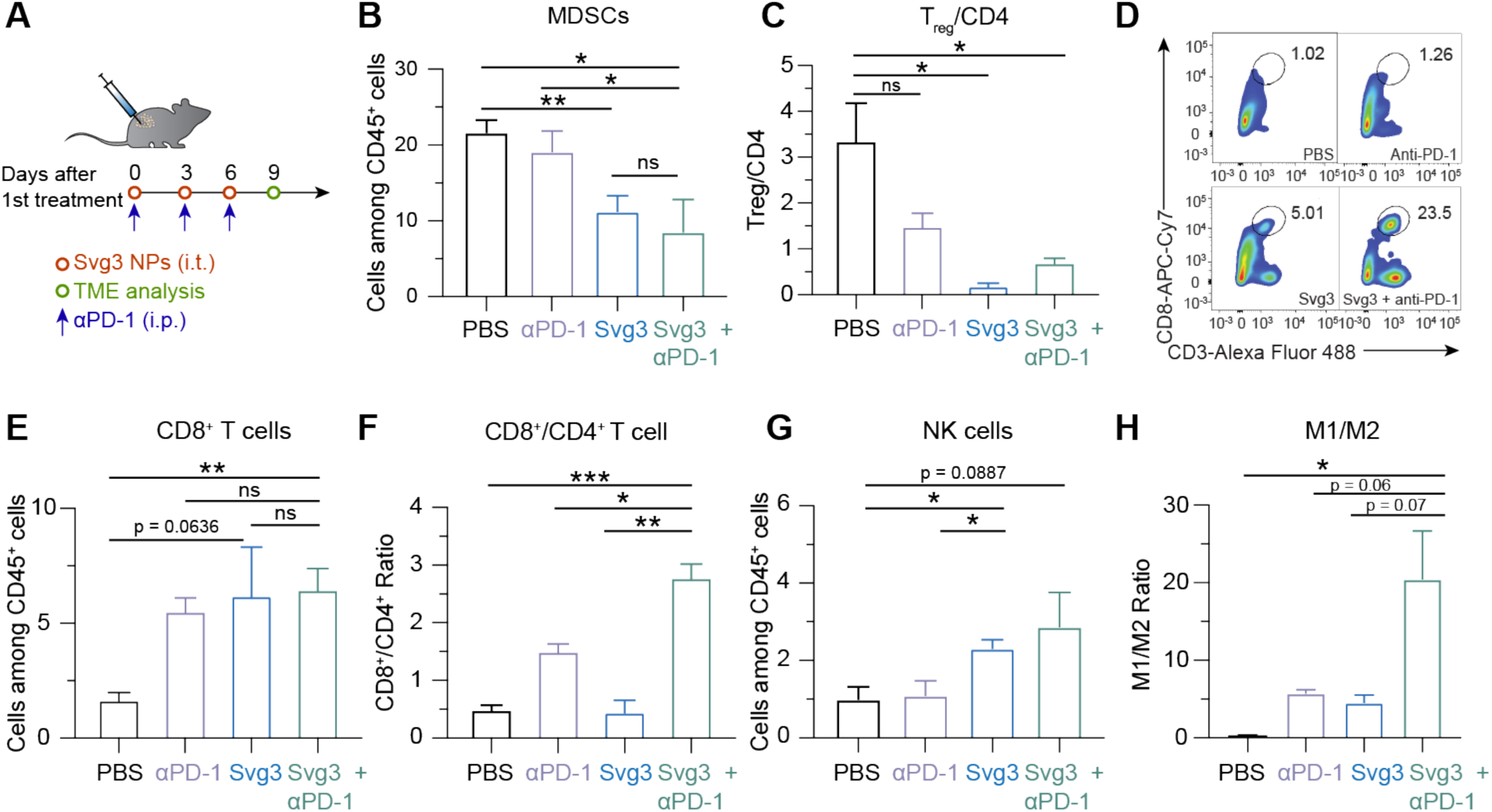
Liposomal Svg3 reduced TME immunosuppression. **A**) Timeline of TME immune cell subset analysis in a 4T1 tumor xenograft model treated with liposomal Svg3 (i.t.), alone or combined with αPD-1 (i.p.) as shown above. Tumors were s.c. injected on the flank of Balb/c mice (*n* = 3-5). Tumors were harvested on day 9 for single cell analysis by flow cytometry. **B**) Percentage of MDSCs among CD45^+^ cells in as-treated 4T1 tumors. **C**) Ratio of CD4^+^Foxp3^+^CD25^+^ Treg over all CD4^+^ T cells in as-treated 4T1 tumor. **D**) Representative flow cytometric graphs of CD8^+^CD3^+^ T cells in as-treated 4T1 tumors. **E**) Percentage of CD8^+^ T cells among CD45^+^ cells in as-treated 4T1 tumors. **F**) Ratio of CD8^+^ T cells over CD4^+^ T cells in 4T1 tumors. **G**) Densities of NK cells among CD45^+^ cells in as-treated 4T1 tumors. **H**) Ratio of M1-like macrophages over M2-like macrophages in as-treated 4T1 tumors. Data: mean ± S.E.M. *P* values were determined by t test (ns: not significant, *p* > 0.05; \**p* < 0.05; \*\**p* < 0.01; \*\*\**p* < 0.001; \*\*\*\**p* < 0.0001).

### Combination of Svg3 and αPD-1 for robust tumor immunotherapy

The abilities of Svg3 to elicit IFN-I responses and reduce TME immunosuppression make this immunostimulant appealing for ICB combination immunotherapy of cancer. Specifically, we test this for Svg3 as both a therapeutic immunostimulant and an adjuvant for cancer antigen vaccines. We first evaluated Svg3 as an immunostimulant for the ICB combination immunotherapy of multiple poorly immunogenic murine tumor models in syngeneic mice, including 4T1 mammary carcinoma in Balb/c mice, and B16 melanoma and MOC2 oral squamous cell carcinoma (OSCC) in C57BL/6 mice. Worth noting, upon Svg3 treatment, the IFN-β production levels by these tumor cells varied significantly (**Supplementary Fig. 5)**. To investigate the therapeutic efficacy of Svg3 in vivo, B16F10 tumor cells were s.c. inoculated in the right flank of syngeneic C57BL/6 mice. When tumors were approximately 50 mm^3^, mice started to be treated with liposomal Svg3 (i.t., 5 x 3 nmole) and αPD-1 (i.p., 5 x 100 μg) every three days for five times. While liposomal Svg3 monotherapy inhibited tumor progression, especially in B16F10 tumor, combination of liposomal Svg3 with αPD-1 dramatically enhanced the tumor therapeutic efficacy in all these tumor models (**Fig. 6B-F**). Consistently, liposomal Svg3 + αPD-1 significantly prolonged mouse survivals (**Fig. 6E**). Worth noting, none of these treatments caused any significant reduction of mouse body weights (**Supplementary Fig. 6**), suggesting their potential of great safety (**Fig. 6G**). Meanwhile, in naïve C57BL/6 mice, we studied the systemic innate immune responses elicited by Svg3 by measuring a panel of serum chemokines and cytokines by Luminex. As a result, a single dose of s.c. administered liposomal Svg3 significantly elevated the serum levels of proinflammatory TNF-α and T cell recruiting chemokine CXCL-10, but not IL-6. (**Fig. 6H**). The increased expression of anti-inflammatory cytokine or chemokine IL-13 and eotaxin may also indicate rapid activation of CD4^+^ T cells upon exposure to Svg3.

**Figure 6.**
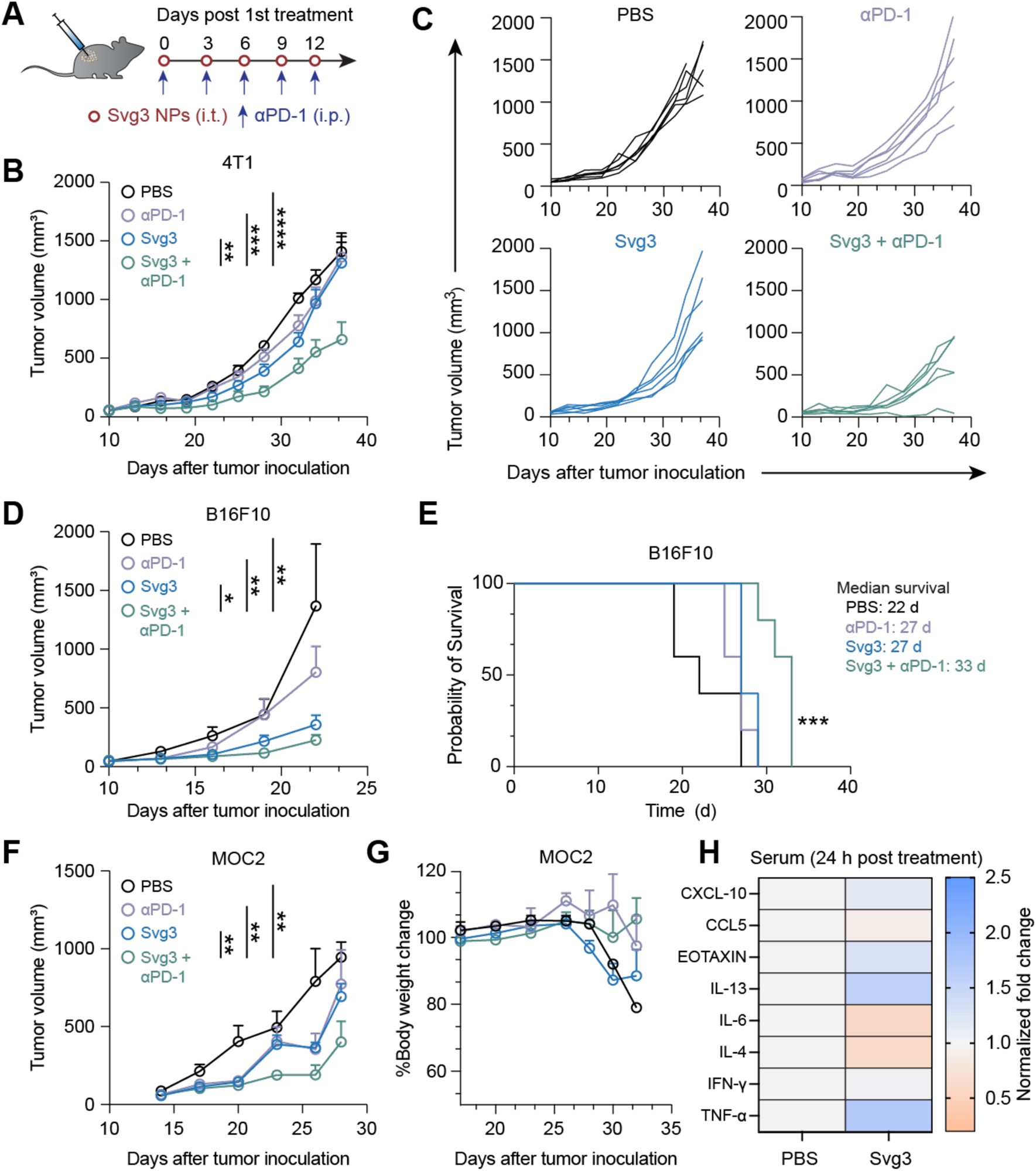
Svg3 improved ICB therapeutic efficacy in multiple poorly immunogenic tumor models. **A**) Timeline of tumor therapy studies in mice. When tumors were approximately 50 mm^3^, mice were treated with liposomal Svg3 (i.t., 5 x 3 nmole) and αPD-1 (i.p., 5 x 100 μg) every three days for five times. **B**) Average 4T1 tumor growth curves. **C**) Individual 4T1 tumor growth curves. **D**) Average B16F10 tumor growth curves. **E**) Meier-Kaplan survival curves of B16F10 tumor-bearing mice after the above treatments. **F**) Average MOC2 tumor growth curves. **G**) Body weights of MOC2 tumor-bearing mice after the above treatments. **H**) Luminex analysis of serum cytokines and chemokines 24 h after mice were s.c. treated with liposomal Svg3. Data: mean ± S.E.M. p values were determined by two-way ANOVA (\*\**p* < 0.01; \*\*\**p* < 0.001; \*\*\*\**p* < 0.0001).

We further investigated the roles of cGAS and STING in Svg3-mediated anti-tumor therapy using cGAS KO mice [B6(C)-*Cgas^tm^*^1d^(EUCOMM)*^Hmgu^*/J or *cGas*^-/-^] and Goldenticket STING KO mice (C57BL/6J-*Sting1^gt^*/J or *Sting*^gt/gt^), respectively. Mice with s.c. B16F10 tumors (50 mm^3^) were again treated as above with five doses of liposomal Svg3, with liposomal STING agonist 2’3’-cGAMP as a control, alone or combined with αPD-1. In *cGas*^-/-^ mice, B16F10 tumors, Svg3 lost its ability to improve the tumor therapeutic efficacy of αPD-1, suggesting that cGAS is essential for the tumor therapeutic efficacy of Svg3 (**Fig. 7A**). In contrast, 2’3’-cGAMP significantly enhanced the ability of αPD-1 to retard tumor progression (**Fig. 7A**), verifying the intact STING and downstream immunostimulatory signal pathway in this mouse model. Moreover, in *Sting*^gt/gt^ mice, neither Svg3 nor 2’3’-cGAMP significantly potentiated the tumor therapeutic efficacy of αPD-1 (**Fig. 7B**). This suggests that STING is essential for the tumor therapeutic efficacy of Svg3. Collectively, these results demonstrated the cGAS-STING-specific activation by Svg3 to potentiate the tumor therapeutic efficacy of ICB for combination immunotherapy.

**Figure 7.**
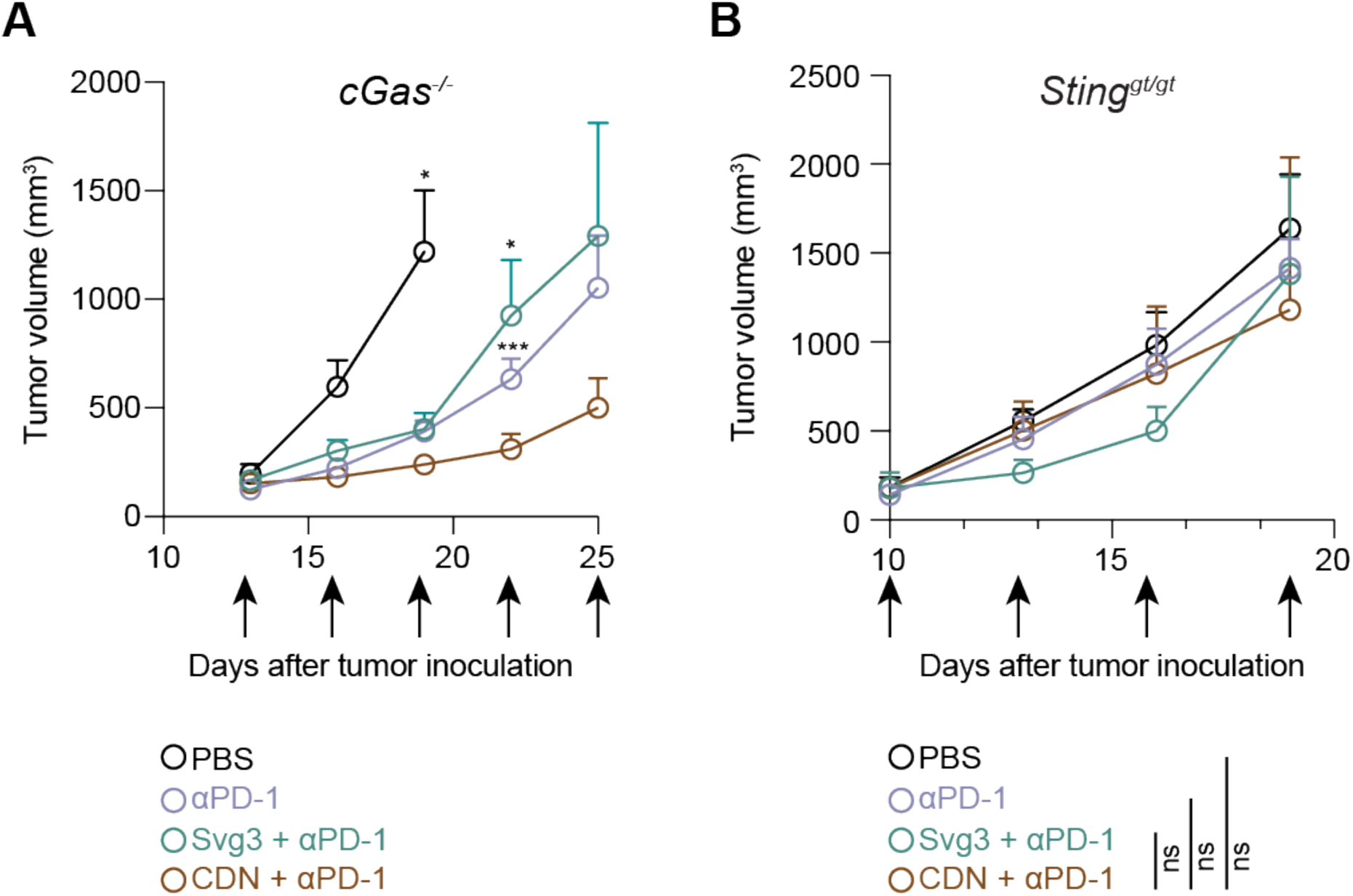
cGAS and STING are essential for Svg3 to potentiate ICB tumor immunotherapy in mice. **A**) B16F10 tumor growth curves in *cGas*^-/-^ mice treated with liposomal Svg3, with liposomal 2’3’-cGAMP as a control, alone or combined with αPD-1. Svg3, but not 2’3’-cGAMP, lost the ability to potentiate the tumor therapeutic efficacy of αPD-1. **B**) B16F10 tumor grow curves in *Sting*^gt/gt^ mice treated as above. Neither Svg3 nor 2’3’-cGAMP significantly potentiated the tumor therapeutic efficacy of αPD-1. Arrows mark treatments. Liposomal Svg3 and 2’3’-cGAMP: i.t., 5 x 3 nmole; αPD-1: i.p., 5 x 100 μg. Data: mean ± S.E.M. P values were determined by *t*-test (A) at day 19 (2’3’-cGAMP + αPD-1 vs, PBS) or at day 22 (αPD-1 or Svg3 + αPD-1 vs. 2’3’-cGAMP + αPD-1), or by two-way ANOVA (B) (ns: not significant, *p* > 0.05; \**p* < 0.05, \*\*\**p* < 0.001).

### As a vaccine adjuvant, Svg3 promoted antigen presentation and antigen-specific T cell responses

Because IFN-I responses promote antigen presentation^[25]^, we also evaluated Svg3 as an immunostimulatory adjuvant of peptide subunit vaccines to promote antigen presentation and T cell priming. The three essential signals for APCs to present antigens to T cells include co-stimulatory factors, proinflammatory cytokines, as well as the cell surface presentation of MHC-epitope complexes. In murine DC2.4 cells, Svg3 upregulated the expression of costimulatory factors CD80, CD86, and CD40 (**Fig. 8A**), and enhanced the production of proinflammatory cytokines IFN-β and IL-6. Next, we studied antigen presentation using a model antigen chicken ovalbumin (OVA), which has an MHC-I-restricted epitope OVA_257-264_ (SIINFEKL). We treated DC2.4 cells with admixed Svg3 and OVA for 24 h, followed by immunostaining of H-2K^b^/SIINFEKL complexes using a dye-labeled antibody (H-2K^b^ is a haplotype of murine MHC-I). Flow cytometric analysis showed that Svg3 enhanced SIINFEKL presentation on DC2.4 cells at a level comparable to that by CpG, a benchmark potent Toll-like receptor 9 (TLR9) agonist (**Fig. 8B**). We expect that Svg3 would bypass the limitations associated with the clinical translation of CpG, such as limited TLR9-positive immune cell subsets, especially in human, as well as the inter-species discrepancy of TLR9^[26]^. To study the impact of Svg3 on SIINFEKL-specific T cell priming, we used immortalized SIINFEKL-specific B3Z CD8^+^ T cell hybridoma that, upon recognition of H-2K^b^/SIINFEKL complexes on DCs, are activated to produce β-galactosidase (β-gal) ^[27]^. DC 2.4 cells were first treated with Svg3 + OVA (24 h), and the treated DCs were co-cultured with B3Z CD8^+^ T cells. As shown by β-gal activity, Svg3 + OVA elicited 5-fold B3Z activation relative to OVA alone (**Fig.8C**), which demonstrated the ability of Svg3 as a vaccine adjuvant to promote T cell priming. These data demonstrated the potential of Svg3 to promote antigen presentation and T cell responses.

**Figure 8.**
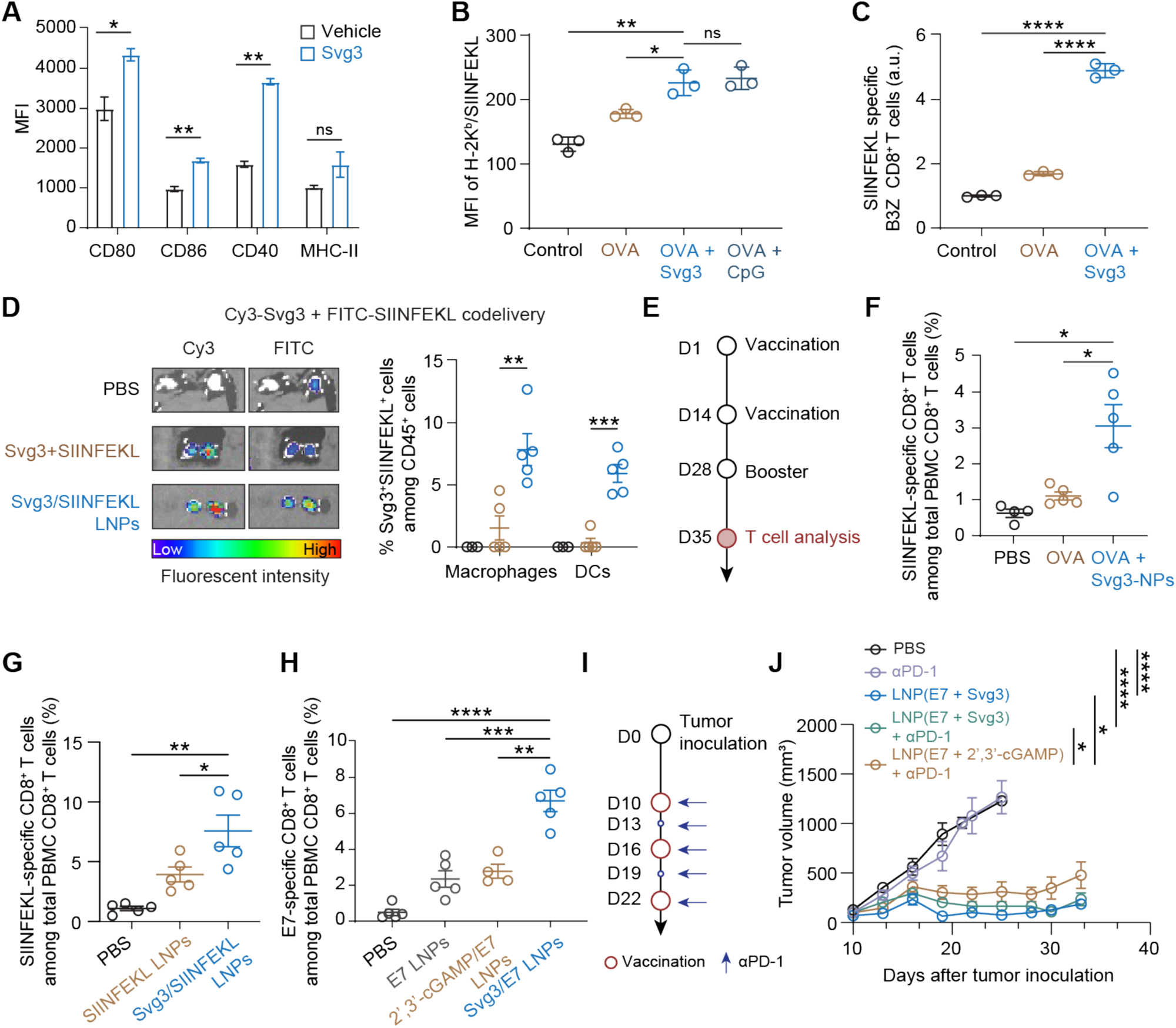
As a vaccine adjuvant, Svg3 potentiated the immunogenicity of poorly immunogenic peptide antigens to elicit robust T cell response for potent tumor combination immunotherapy. **A**) Flow cytometry results showing that Svg3 elevated the expression of costimulatory factors (CD80, CD86, and CD40) as well as MHC-II on RAW 264.7 macrophages. Cells were transfected with Svg3 LNPs for 24 h. **B**) Flow cytometric quantification of the H-2K^b^/SIINFEKL complexes on as-treated DC2.4 cells showed that Svg3 promoted SIINFEKL presentation. Cells were transfected with vaccine-loaded LNPs for 24 h. CpG was used as a control. MFI: mean fluorescence intensity. **C**) The β-gal activity in B3Z T cell hybridoma cocultured with vaccine-treated DC2.4 cells, suggesting that Svg3 treatment of DC2.4 cells enhanced the priming of B3Z T cells. Vaccine treatment in DC2.4 cells: 24 h; coculturing of DC2.4 cells and B3Z cells: 24 h. β-gal activity was measured by the 570 nm absorbance upon β-gal-catalyzed substrate conversion. **D**) Left: fluorescence imaging of ex vivo inguinal lymph nodes from C57BL/6 mice 16 h after s.c. injection (at tail base) of the indicated Cy3-Svg3 + SIINFEK_(FITC)_L formulations or PBS. Right: as quantified from flow cytometric analysis, the fraction of Svg3^+^SIINFEK_(FITC)_L^+^ APC subsets among CD45^+^ cells in inguinal lymph nodes, suggesting that LNPs promoted the codelivery of Svg3 and SIINFEKL into intranodal APCs. **E**) Immunization study timeline of Svg3-adjuvanted peptide vaccines. Immunization: s.c. at mouse tail base. Svg3 and peptides were delivered by LNPs, OVA was injected as free solutions or mixed with Svg3-LNPs (doses: 3 nmol Svg3, 25 μg peptides or proteins). **F-G**) Tetramer staining results showing the percentages of SIINFEKL-specific CD8^+^ T cells among total PBMC CD8^+^ T cells in mice immunized with OVA protein vaccines (F) (day 35) and SIINFEKL peptide vaccines (G) (day 21), respectively. **H**) Tetramer staining results showing the percentages of E7-specific CD8^+^ T cells among total PBMC CD8^+^ T cells (day 35) in mice as immunized above (days 0, 14, 28). 2’3’-cGAMP was used as a control. I) Study timeline of the combination immunotherapy therapy of E7-positive TC-1 tumor in C57BL/6 mice. **J**) TC-1 tumor growth curves. Treatments were initiated when tumors reached ∼100 mm^3^. LNP vaccines: s.c. at tail base, 3 nmole Svg3 or 2’3’-cGAMP, 25 ug E7 peptide; αPD-1: i.p., 5 x 100 μg. Data: mean ± S.D. (A-C), and mean ± S.E.M (D-J). *P* values were determined by t-test (A-H) or two-way ANOVA (J) (ns: not significant, *p* > 0.05; \**p* < 0.05, \*\**p* < 0.01; \*\*\**p* < 0.001; \*\*\*\**p* < 0.0001).

We then studied Svg3 as an adjuvant for peptide vaccines to elicit T cell responses in C57BL/6 mice. Svg3 and SIINFEKL peptide were co-loaded in ionizable lipid DLin-MC3-DMA-based LNPs, with a >95% of Svg3 encapsulation efficiency, *ca*. 170 nm in diameter, and nearly neutral electrostatic charge (**Supplementary Fig. 2A**). LNPs dramatically enhanced the intracellular delivery of Svg3 in RAW 264.7 cells (**Supplementary Fig. 2D**). Svg3/SIINFEKL LNPs were s.c. administered at mouse tail base, allowing efficient accumulation of Svg3 and SIINFEKL in draining inguinal lymph nodes (**Fig. 8D**). Flow cytometric analysis of intranodal single cell subsets further revealed the efficient uptake of LNPs by key APC subsets, including DCs and macrophages (**Fig. 8D**). Such efficient co-delivery of antigen and Svg3 adjuvant to APCs is critical for the optimal T cell responses with minimal immune tolerance caused by unadjuvanted peptide antigen exposure in APCs. We further studied Svg3 as an adjuvant to potentiate the immunogenicity of protein and peptide antigens to elicit antigen-specific CD8^+^ T cells, using nanocarriers co-loaded with Svg3 and antigens. We first used two model antigens, chicken ovalbumin (OVA) and an MHC-I-restricted peptide antigen OVA_257-264_ (SIINFEKL). As shown by a tetramer staining assay, relative to OVA alone, Svg3-adjuvanted OVA (*s.c*. injection at tail base on days 0, 14, 28) elicited a significantly high fraction of SIINFEKL-specific CD8^+^ T cells among peripheral blood mononuclear cells (PBMCs) (**Fig. 8F**). Consistently, relative to peptide SIINFEKL-loaded LNPs, Svg3/SIINFEKL-co-delivering LNPs (s.c. injection at tail base, days 0, 14) elicited 2-fold SIINFEKL-specific CD8^+^ T cells among total live PBMC CD8^+^ T cells (day 21) (**Fig. 8G**). These results demonstrated the ability of Svg3 to potentiate the immunogenicity of antigens to elicit T cell responses.

We then studied Svg3 adjuvant to elicit CD8^+^ T cell responses against an MHC-I-restricted E7 peptide, an oncoviral antigen derived from E7 protein in human papillomavirus (HPV) 16 and 18. HPV infection causes various types of cancer, and high-risk HPV (e.g., HPV16) causes ∼5% of all cancer cases^[28]^. Therapeutic vaccines hold the potential to treat these cancers with defined oncoviral antigens. However, current preventive HPV vaccines and E7-based experimental therapeutic vaccines have limited efficacy against HPV-associated tumors in the clinic, in part due to low immunogenicity of HPV antigens.^[28–30]^ This demands novel technologies such as potent and broadly applicable adjuvants to potentiate the immunogenicity of HPV vaccines. To this end, Svg3 and E7 peptide were coloaded in MC3 LNPs, with STING agonist 2’3’-cGAMP as a control for Svg3. In C57BL/6 mice, immunization of the resulting Svg3/E7 LNPs (s.c. at tail base, days 0, 14, 28) significantly enhanced the fraction of E7-specific CD8^+^ T cells among total PBMC CD8^+^ T cells (**Fig. 8H**), which outperformed cGAMP/E7-loaded LNPs. Giving the central role of CD8^+^ T cells in tumor immunotherapy, Svg3/E7 LNPs were then evaluated for tumor combination immunotherapy of E7-positive TC-1 murine tumor (∼100 mm^3^) in syngeneic C57BL/6 mice (**Fig 8I**). While Svg3/E7 LNPs alone already showed significant tumor inhibition, combining Svg3/E7 LNPs with αPD-1 further enhanced the tumor therapeutic efficacy, resulting in complete regression of 2 out of 7 tumors (**Fig. 8J**). These results clearly demonstrate the potential of Svg3 as a vaccine adjuvant for cancer combination immunotherapy.

## Conclusion

Cancer immunotherapy, which leverages the host immune system to treat cancer, has significantly improved the treatment outcomes for many types of cancer. ICB is one of the current mainstream approaches to cancer immunotherapy. However, most of cancer patients do not benefit from current ICB, due to a lack of pre-existing antitumor immune cells and immune checkpoints, as well as TME immunosuppression, among others. Moreover, some patients suffer from immune-related adverse events caused by the imbalanced immune homeostasis. This demands innovative approaches, such as cancer therapeutic vaccines, for combination immunotherapy with ICB to maximize the therapeutic potential of ICB. Cancer therapeutic vaccines can reduce systemic and tumor immunosuppression, elicit and augment antitumor innate and adaptive immunity, upregulate the expression of immune checkpoints and thereby sensitizing them for ICB, and promote the tumor infiltration of antitumor cells and molecules. Immunostimulants, including PRR agonists, are extensively used in therapeutic vaccines as either non-tumor-specific innate immunostimulants or adjuvants for tumor-specific antigens. cGAS-STING immunostimulation pathway plays a critical role in innate immunity that mediate anti-cancer therapies, including ICB. Upon dsDNA binding, cGAS is activated to synthesize cGAMP, which then stimulates STING to trigger anti-tumor IFN-I responses. Current approaches to the development of cancer therapeutic cGAS-STING-activating immunostimulants have been almost exclusively targeting STING. However, current STING agonists have shown limited tumor therapeutic efficacy in the clinic, largely due to their hydrophilicity, often negative electrostatic charges and susceptibility to enzymatic degradation, which results in poor bioavailability. For example, i.t. administration of a CDN, MK-1454, showed minimal therapeutic efficacy in patients with solid tumors or lymphomas ^[31]^. In addition, a Phase I clinical trial data of i.t. administered STING agonist ADU-S100 in combination with αPD-1 (spartalizumab) showed modest clinical benefit ^[17]^.

Compared with current small molecule-based STING agonists, cGAS agonistic oligonucleotide therapeutics may have the following advantages: 1) reproducible and economic production using existing automated cGMP manufacturing facilities, 2) advanced drug delivery systems that dramatically improve the pharmacokinetics and therapeutic efficacy of oligonucleotides, 3) well-established oligonucleotide chemistry that may further improve the biostability, pharmacokinetics, safety, and therapeutic efficacy of oligonucleotide therapeutics, and 4) cGAS agonists can bypass the complications associated with the allele selectivity of STING in human. Here, by structural optimization, we developed a cGAS-agonistic oligonucleotide, Svg3, for versatile applications in ICB combination cancer immunotherapy. Svg3 specifically activated cGAS, eliciting potent IFN-I responses not only in murine cells but also in human cells and tissues, which are pivotal for its future clinical translation. Remarkably, Svg3 outperformed several state-of-the-art STING agonists to elicit IFN-I responses in murine and human immune cells. Moreover, Svg3 did not significantly elevate AIM2-associated *Il18* or *Il1b* expression or elicit detectable IL-18 or IL-1β production in RAW 264.7 macrophages. This rules out the possibility for Svg3 to activate AIM2, another cytosolic dsDNA sensor that can induce confounding inflammasome activation in these cells.

Svg3 was easily formulated in well-established liposomes and LNPs, which improved the intracellular delivery, endosome escape, as well as tissue retention of Svg3 in tumors (for i.t. Svg3 administration) and draining lymph nodes (for s.c. Svg3 administration). I.t. Svg3 nanoparticles administration reduced multi-faceted immunosuppression and enhanced antitumor immunity in tumor. As a result, Svg3 significantly improved the therapeutic efficacy of ICB in multiple syngeneic tumor models, in wildtype mice but neither *cGas^-^*^/-^ mice nor *Sting*^gt/gt^ mice. Further, as an adjuvant, Svg3 was codelivered with peptide antigens in LNPs to draining lymph nodes. As a result, Svg3 potentiated the immunogenicity of the otherwise poorly immunogenic peptide antigens to elicit T cell responses in mice, resulting in robust immunotherapy of cancer.

Further comprehensive optimization of Svg3 may further improve its cGAS activation efficacy. Future structural and biochemical studies of cGAS binding and activation by Svg3 may reveal the underlying mechanism of action and provide insight for its preclinical and clinical development. Moreover, future studies will explore systemic delivery of Svg3 for the treatment of surgically inaccessible tumors and metastatic tumors, in which i.t. administration may find limited applicability. Systemic delivery of immunostimulants would likely elevate the systemic cytokine levels, which may lead to immune toxicity in the scenario such as cytokine storm. Therefore, it will be important to optimize Svg3 formulation to have a balanced efficacy and safety. To this end, recent research has demonstrated the feasibility of systemic delivery of vaccines, such as STING agonists and TLR7/8 agonist-adjuvanted nanovaccines^[32,33]^. Overall, oligonucleotide-based cGAS agonists represent a promising approach as immunostimulant therapeutics and vaccine adjuvants for cancer combination immunotherapy.

## Methods and Materials

### Cell culture

DC2.4 cells, 4T1 cells, and TC-1 were cultured in RPMI 1640 medium. RAW-ISG cells were obtained from InvivoGen and cultured in DMEM medium containing 100 µg/mL Normocin and 200 µg/mL Zeocin. MOC2 cells were cultured in DMEM/F12 medium. RAW 264.7 cells and B16F10 cells were cultured in DMEM medium. B3Z cells and THP-1 cells were cultured with RPMI 1640 medium supplemented with 2 mM L-glutamine, 1 mM sodium pyruvate, and 50 μM 2-mercaptoethanol. All medium was supplemented with 10% FBS and 0.1% penicillin and streptomycin. All cells were cultured in a humidified atmosphere (5% CO_2_, 37 °C) in a Biosafety Level II incubator.

### IFN-I expression

IFN-I cytokine levels were measured by ELISA in RAW264.7 cells and by using a IFN reporter cell in THP-1 cells. RAW264.7 cells were seeded in 96-well plate at density of 5X10^4^ cells/well and incubated overnight for experiment. Svg3 was transfected into cells using lipofectamine reagent at a final concentration of 25 nM and incubated at 37 ℃ and 5% CO_2_ for 24 h. The medium was collected to test the concentration of IFN-β. THP-1 cells were seeded at densities of 1×10^4^ cells/well in a 96-well plate. Cells were treated with Svg3 or cGAMP respectively for 24 h. At the same time, HEK-Blue™ IFN-α/β cells (Invivogen) were seeded at densities of 1×10^4^ cells/well in a 96-well plate. Then THP-1 cell supernatant was added to HEK-Blue™ IFNα/β cells for 24 h. 50 μL cell supernatant was collected from each sample and added to 150 μL of QUANTI-Blue SEAP detection medium (InvivoGen) and incubated for 2 h at 37 °C. SEAP activity was assessed by measuring the absorbance at 630 nm on a plate reader.

### ISG expression

RAW-Lucia^TM^ ISG cells, which were generated from RAW264.7 cell line by stable integration of an IRF-inducible Lucia luciferase reporter construct, were used to evaluate cGAS activation. To validate the role of cGAS in Svg3 stimulation, cGAS knock-out RAW-Lucia^TM^ ISG cells were used at the same Svg3 treatment condition. For comparison of Svg3 and STING agonists, Svg3 and STING agonists including 2’3’-cGAMP, ADU-S100 or diABZI were transfected at different concentrations respectively, using lipofectamine reagent. The expression of IRF-induced Lucia luciferase in the culture medium was measured using QUANTI-Luc^TM^ following the manufacturer’s instruction.

### 2’3’-cGAMP production by Svg3-activated cGAS

RAW 264.7 macrophages were seeded in 12-well plates and incubated overnight. Svg3 or ISD was transfected using lipofectamine 2000 at a final concentration of 100 nM. The cells were collected at different time points after transfection, and lysed using RIPA lysis buffer containing protease inhibitors and EDTA on ice for 15 min. The concentrations of 2’3’-cGAMP was measured using 2’3’-cyclic cGAMP ELISA kit (Invitrogen) following the manufacturer’s instruction.

### Svg3 binding affinity with cGAS

Cy5-labeled Svg3 (Cy5-Svg3) was used to measure the binding affinity with cGAS by MST, as described previously^[34]^. In brief, the concentration of Cy5-Svg3 was determined to ensure the fluorescence intensity was between 800-1000. cGAS was diluted using Tris buffer containing 0.05% Tween 20. Cy5-Svg3 of a series of different final concentrations was added into cGAS solutions and incubated for 15 min at room temperature. Then 20 μL sample was loaded into standard treated capillaries for measurements on a Monolith NT Automated (Nanotemper).

### Co-stimulatory factor expression

RAW 264.7 macrophages and DC2.4 DCs were respectively seeded in 12-well plate and incubated overnight for experiment. Svg3 was transfected using lipofectamine reagent at a final concentration of 25 nM. After treatment for 24 h, the cells were collected and stained with anti-mouse CD86-PerCp, anti-mouse CD80-Alexa Fluor 647, anti-mouse CD40-FITC, anti-mouse I-A/I-E (MHC II)-PE respectively for flow cytometry (BD Canto).

### B3Z T cell activation

B3Z is a SIINFEKL-specific CD8^+^ T cell hybridoma that can be activated by recognizing SIINFEKL/H-2K^b^ complex^[27]^. Activated B3Z cells produce β-galactosidase (β-gal), which can hydrolyze the substrate of chorophenol red-β-D-galactopyranoside (CPRG) into red product. To evaluate B3Z activation, DC2.4 cells were seeded in 12-well plate and treated with lipo-transfected Svg3 with ovalbumin or free ovalbumin for 24 h. After incubation, the medium was removed and B3Z cells were cocultured for another 24 h. Then cells were lysed for 4 h at 37 °C with lysis buffer (PBS with 100 mM 2-mercaptoethanol, 9 mM MgCl_2_, 0.2% Triton X-100 and 0.15 mM CPRG). The reaction was stopped by 1 M sodium carbonate. The magnitude of antigen priming was evaluated through absorbance measurements (λ = 570 nm).

### RNA-seq transcriptomic analysis of Svg3-treated mouse macrophage

RAW 264.7 macrophages were seeded in 6-well plate and treated with lipo-transfected Svg3 at a final concentration of 25 nM. The control group was lipofectamine without Svg3. After 6 h incubation, the culture medium was removed, and the cells were collected for RNA extraction and sequencing. An average of 30 million paired end reads of length 150 base pair (bp) were generated for each sample. The quality of RNA-Seq reads was assessed with FastQC v0.11.9. The reads were aligned using STAR aligner^[35]^ version 2.7.6a to reference genome GRCm38. Raw gene counts of mapped reads were aggregated using featureCounts^[36]^. The differential gene expression analysis was performed with Bioconductor package DESeq2 v1.30.0 ^[37]^ using the normalized and filtered counts per gene from the RNA-Seq data.

### Quantitative PCR

RAW 264.7 macrophages were seeded in 12-well plates and transfected with DNA at a final concentration of 25 nM. After incubation for 24 h, the cells were collected, and the RNA were extracted using RNA extraction kit (Invitrogen) following manufacturer’s protocol. RNA was reverse-transcribed to cDNA using High-Capacity cDNA Reverse Transcription Kit (Thermo Fisher). qPCR was carried out using SYBR Green Master Mix (Applied Biosystems) on a QuantStudio 3 system (Applied Biosystems).

### IFN responses in human tumor tissue

Human head and neck cancer tissues were provided by VCU Massey Cancer Center Tissue and Data Acquisition and Analysis Core. Fresh tumor tissues were evenly cut into cubes and randomly assigned into groups. The resulting tissues were cultured in 1 mL complete DMEM medium in 6-well plates. 0.3 nmol Svg3 loaded in liposomes was injected into the above tissues using syringe needles. After 24 h incubation in a tissue culture incubator, the tumor tissue RNA was extracted using RNA extraction kit (Invitrogen). RNA was reverse-transcribed to cDNA using High-Capacity cDNA Reverse Transcription Kit (Thermo Fisher). qPCR was carried out using SYBR Green Master Mix (Applied Biosystems) on a QuantStudio 3 system (Applied Biosystems).

### In vitro antigen presentation

DC2.4 cells were seeded in 6-well plate and incubated with lipo-transfected Svg3 with ovalbumin or free ovalbumin for 24 h. Treated DC 2.4 cells were collected, washed with DPBS, and stained with allophycocyanin (APC)-conjugated anti-SIINFEKL/H-2K^b^ antibody (BioLegend) for 30 min and then analyzed by flow cytometry (BD Canto).

### Nanoparticle preparation

#### Liposomes

Svg3 was formulated into liposome using thin film hydration method as previously reported^[38]^. In brief, DOTAP, cholesterol, DOPE and DSPE-PEG2000 were dissolved in chloroform and mixed in a round-bottom bottle. The thin lipid film was formed by the removal of chloroform using rotary evaporator. Svg3 aqueous solution then was added and vortexed vigorously for 30 min. The N/P ratio for lipids and DNA was 10. The crude liposomes were extruded using extruder (Avanti). Liposomes were diluted in 1XDPBS and measured using Zetasizer (Malvern Panalytical) for size and zeta potential measurement.

#### LNPs

Svg3 was formulated in lipid nanoparticles using ethanol dilution method. In brief, DLin-MC3-DMA, cholesterol, DOPC and DMG-PEG2000 were dissolved in ethanol and DNA was dissolved in 10 mM citrate buffer (pH 4). All lipids were mixed with a molar ratio of 50: 38.5:10:1.5. The two solutions were rapidly mixed at room temperature for 15 min, at an aqueous to ethanol ratio of 3:1(v/v) with a final mass ratio of lipids and DNA of 20:1. LNPs were diluted in 1X DPBS and measured using Zetasizer (Malvern Panalytical) to determine size and zeta potential. The fresh LNPs were dialyzed using Pur-A-Lyzer^TM^ dialysis kit (Sigma-Aldrich, MWCO: 6 KDa) at 4 °C overnight, and concentrated by ultracentrifuge tubes (Millipore, MWCO: 10K) for in vivo injection.

### Cell uptake of liposomal Svg3

Fluorescent-labeled Svg3 was used to evaluate the cellular uptake after being formulated into liposome using method above. RAW 264.7 cells were seeded in 6-well plate and cultured overnight for further experiment. Cy3-labeled Svg3 liposomes were diluted using serum-free medium and incubated at a final concentration of 100 nM per well. The medium was discarded, and cells were washed with 1XDPBS and then collected for flow cytometry (BD FACSCanto).

### Animal studies

C57BL/6 mice and Balb/c mice (6–8 weeks old) were purchased from Charles River Laboratory. B6(C)-Cgastm1d(EUCOMM)Hmgu/J and C57BL/6J-Sting1gt/J mice were ordered from Jackson Laboratory. All animals were maintained at the animal facilities of Virginia Commonwealth University under specific pathogen-free conditions and treated in accordance with the regulations and guidelines of the Institutional Animal Care and Use Committee. All animal experiments were approved by the Virginia Commonwealth University (VCU) Institutional Animal Care and Use Committee.

### IVIS imaging of Svg3 nanoparticle retention in tumor

For 4T1 tumor accumulation imaging, IR800-labeled Svg3 liposomes were i.t. injected to 4T1 tumor bearing mice. The tumor retention effect of the IR800-Svg3 liposomes in tumor were visualized with the IVIS optical system for 7 days. The excitation wavelength was 780 nm and the emission wavelength was 794 nm. For inguinal lymph node imaging, LNPs coloaded with Cy3 labeled Svg3 and FITC labeled SIINFEKL were injected s.c. at tail base of C57BL/6 mice. After 16h, the mice were sacrificed, and the inguinal lymph nodes were collected. The excitation wavelength and the emission wavelength for Cy3-Svg3 was 548 nm and 570 nm respectively, and 488 nm and 520 nm for FITC-SIINFEKL.

### Antigen-specific T cell response in mice

C57BL/6 mice were immunized *via* s.c. injection at tail base with different treatments on days 0 and 14, and boosted at day 28: (1) PBS, (2) E7 LNPs, (3) (Svg3 + E7) LNPs, and (4) (2’3’-cGAMP + E7) LNPs.

For E7-specific CD8^+^ T cell analysis, flow cytometry analysis was performed on day 35, peripheral blood cells were collected, and red blood cells were lysed using ACK lysis buffer (Thermo Fisher Scientific) for 5 min, and then the cell pellets were collected by centrifugation. Cells were washed twice in 1X DPBS and stained using Zombie Aqua (Biolegend). Then cells were suspended in cold PBS supplemented with 0.1% FBS and stained with dye-labeled staining cocktail including CD8α-APC/Cy7, CD44-Alexa Fluor 647, CD62L-FITC, PD-1-Brilliant Violet 421 (BioLegend), and H-2D(b) HPV16 E7 49-57 RAHYNIVTF PE-Labeled Tetramer (NIH Tetramer Core Facility). Then cells were washed, resuspended in Cytofix (BioLegend) for 20 min at 4°C. After fixation, cells were washed with Perm/Wash buffer twice and resuspended for flow cytometry (BD LSRFortessa-X20).

For SIINFEKL-specific CD8^+^ T cell analysis, the same staining protocol as above was used except for PE-Tetramer H-2K^b^-restricted SIINFEKL (NIH Tetramer Core Facility) was used. The SIINFEKL tetramer staining was performed at day 56.

### Systemic cytokine and chemokine expression by Luminex

C57BL/6 mice were *s.c.* injected with Svg3 liposomes (3 nmol Svg3) or blank liposomes at tail base. After 24 h, blood was collected and centrifuged for 5 min at 2000 rpm to collect serum. Serum cytokines and chemokines were measured by Luminex at University of Virginia Flow Cytometry Core.

### Tumor therapy

Tumor therapy using i.t. administrated liposomal Svg3 was studied in three murine syngeneic tumor models: 4T1 mammary carcinoma in Balb/c mice, B16F10 melanoma and MOC2 oral cancer in C57BL/6 mice, as well as B16F10 melanoma in *cGas^-^*^/-^ mice [B6(C)-*Cgas^tm^*^1d^(EUCOMM)*^Hmgu^*/J] and Goldenticket *Sting*^gt/gt^ mice (C57BL/6J-*Sting1^gt^*/J) mice. Tumors were respectively established by s.c. injections of tumor cells (5 × 10^5^) into mouse right flank. Tumor volumes and body weights were monitored. Treatment was initiated when the average tumor volumes reached ∼50mm^3^. Liposomal Svg3 was i.t. injected at dose of 3 nmol/mouse every 3 days for 5 times. For ICB, αPD-1 was intraperitoneally injected every 3 days for 5 times. Tumor therapy using Svg3-adjuvanted E7 vaccines was studied in TC-1 tumor in syngeneic C57BL/6 mice. Treatment was initiated when the average tumor size reached 100 mm^3^. Svg3/E7 LNPs and control vaccines was injected s.c. at tail base every 6 days for 3 times. For ICB, mice were i.p. injected with αPD-1 every 3 days for 5 times. Mouse humane endpoint was defined when the body weight dropped by 20% or tumor size reached 2000 mm^3^.

### TME immune analysis

4T1 tumors were established s.c. in Balb/c mice, as previously described, and treated i.t. with PBS, αPD-1, liposomal Svg3, and liposomal Svg3 + αPD-1, respectively, at days 0, 3, and 6. On day 9, tumor tissues were harvested and digested with collagenase A and DNase at 37 °C for 40 min. Digestion mixtures were quenched by adding 10% FBS and samples were filtered through 40-μm cell strainers (Falcon). Cells were stained for 20– 30 min in the dark on ice with the conjugated antibodies as follows (BioLegend unless denoted otherwise) following manufacturer’s recommended concentrations. Myeloid panel: CD45-BV421, CD11b-PE-Cy5, CD11c-Alexa Fluor 594, CD86-PE, Ly-6G/Ly-6C-Alexa Fluor 488, F4/80-FITC, CD206-BV605, and lymphoid cell panel: CD45-BV421, CD3-Alexa Fluor 488, CD4-PerCp-Cy5, CD8α-APC-Cy7, NK-1.1-APC, FoxP3-Alexa Fluor 647. Cells were then fixed with BD Cytofix (BD Biosciences), followed by flow cytometric on a BD LSR Fortessa^TM^ flow cytometer (BD Biosciences) and data analysis using FlowJo software (TreeStar) using gating trees shown in **Supplementary Figure S7-9**.

### Statistical analysis

All statistical analyses were performed using GraphPad Prism software, version 7.0.

### Data availability

All the data related with this study are available within the paper or can be obtained from the authors on request.

## Acknowledgements

G.Z. acknowledges funding support from NIH (R01CA266981, R01AI168684, R35GM143014, R21NS114455), DoD CDMRP Breast Cancer Breakthrough Award Level II (BC210931/P1), American Cancer Society Research Scholar Grant (RSG-22-055-01-IBCD), METAvivor Early Career Investigator Award, VCU Commercialization Fund, and University of Michigan startup fund. The content is solely the responsibility of the authors and does not necessarily represent the official views of the National Institutes of Health. Microscopy was performed at the VCU Microscopy Facility, supported in part by funding from NINDS Center Core Grant 5 P30 NS047463 and, in part, by funding from the NCI Cancer Center Support Grant P30 CA016059. Bioinformatics analysis were provided by VCU Massey Cancer Center Bioinformatics Shared Resource; flow cytometry was performed at the VCU Massey Cancer Center Flow Cytometry Shared Resource; and human tumor tissues were provided by the VCU Massey Cancer Center Tissue and Data Acquisition and Analysis Core; these facilities are all supported, in part, with funding from NIH-NCI Cancer Center Support Grant P30 CA016059. We thank NIH Tetramer Core for providing tetramer reagents, and thank Prof. Xiang-Yang Wang at VCU for providing MOC2 cells.

