## Supplementary Info for "Engineering cGAS-agonistic oligonucleotides as therapeutics and vaccine adjuvants for cancer immunotherapy": 230713 Svg3 SI.pdf

### Contents

**Supplementary Figure 1**. Screening for and characterization of Svg3 as a cGAS agonist.

**Supplementary Figure 2.** Liposomes readily loaded Svg3 for efficient delivery and retention of Svg3 in tissues and cells.

**Supplementary Figure 3.** The transcript levels of *cGAS* and *STING* in three types of human cancers.

**Supplementary Figure 4.** Percentages of DC cells among $CD45^+$ cells in 4T1 tumors.

**Supplementary Figure 5.** Svg3 elicited IFN-I responses in murine tumor cells.

**Supplementary Figure 6.** Mouse body weights during tumor therapy.

**Supplementary Figure 7-9.** Flow cytometry gating trees used in this study.

**Supplementary Table 1.** DNA sequences used for oligonucleotide engineering and screening.

**Supplementary Table 2.** Primer sequences used for qPCR.

**Supplementary Table 3.** Peptide sequences.

**Supplementary Table 4.** Details of antibodies used in this study.

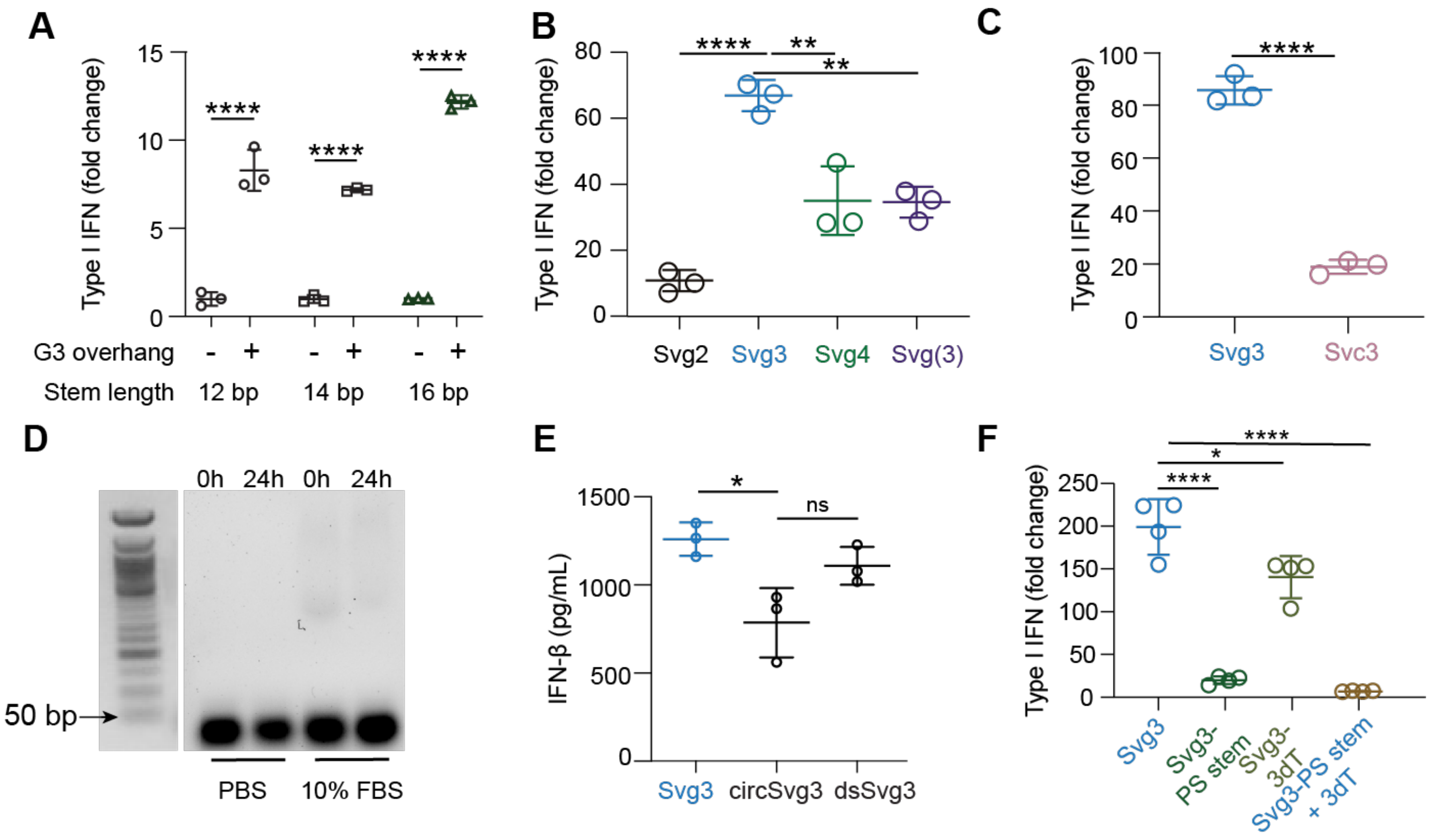

**Supplementary Figure 1**. **Screening for and characterization of Svg3 as a cGAS agonist.** A) IFN-I response in Raw-ISG cells after 24 h transfection with short hairpin DNAs with or without consecutive guanosine in the overhangs. B) IFN-I response in Raw-ISG cells after 24 h transfection with different G content in the overhangs of short hairpin DNAs. C) Comparison of IFN-I response in Raw-ISG cells of hairpin DNA containing consecutive guanosine or cytosine in the overhangs. D) Integrity of Svg3 after incubation in PBS or cell culture medium containing 10% FBS for 24 h verified by 2% agarose gel electrophoresis. E) IFN-β expression in Raw 264.7 cells after transfection of Svg3, circularized Svg3 or double-stranded Svg3 (with two open overhangs) respectively, for 24 h. F) IFN-I response in Raw-ISG cells after 24 h transfection of chemically modified Svg3.

**A**

| | Size/nm | PDI | Zeta potential/mV | EE% Evaluated by gel electrophoresis |
|---|---|---|---|---|
| Svg3 Liposome | 211±5 | 0.15±0.04 | 10.6±0.04 | |
| Svg3 LNP | 179±4 | 0.24±0.04 | 0.97±0.59 | 96.92 ± 0.30 % |
| (Svg3 +E7) LNP | 176±3 | 0.21±0.01 | 1.06±0.66 | NA |
| (cGAMP +E7) LNP | 146±1 | 0.19±0.01 | 0.17±0.23 | NA |

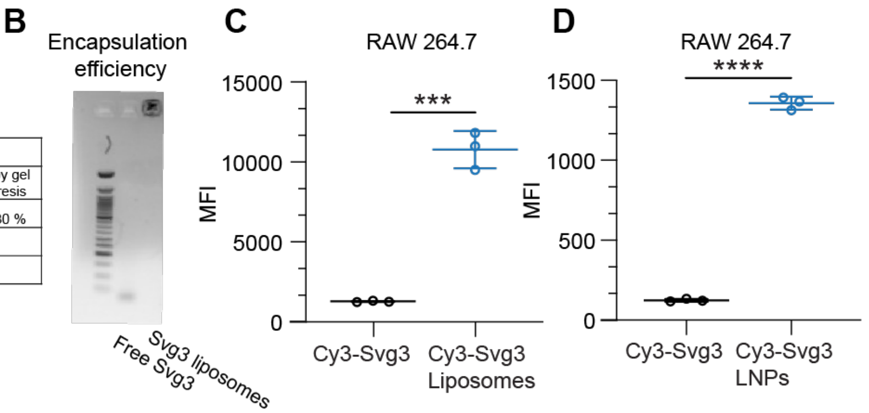

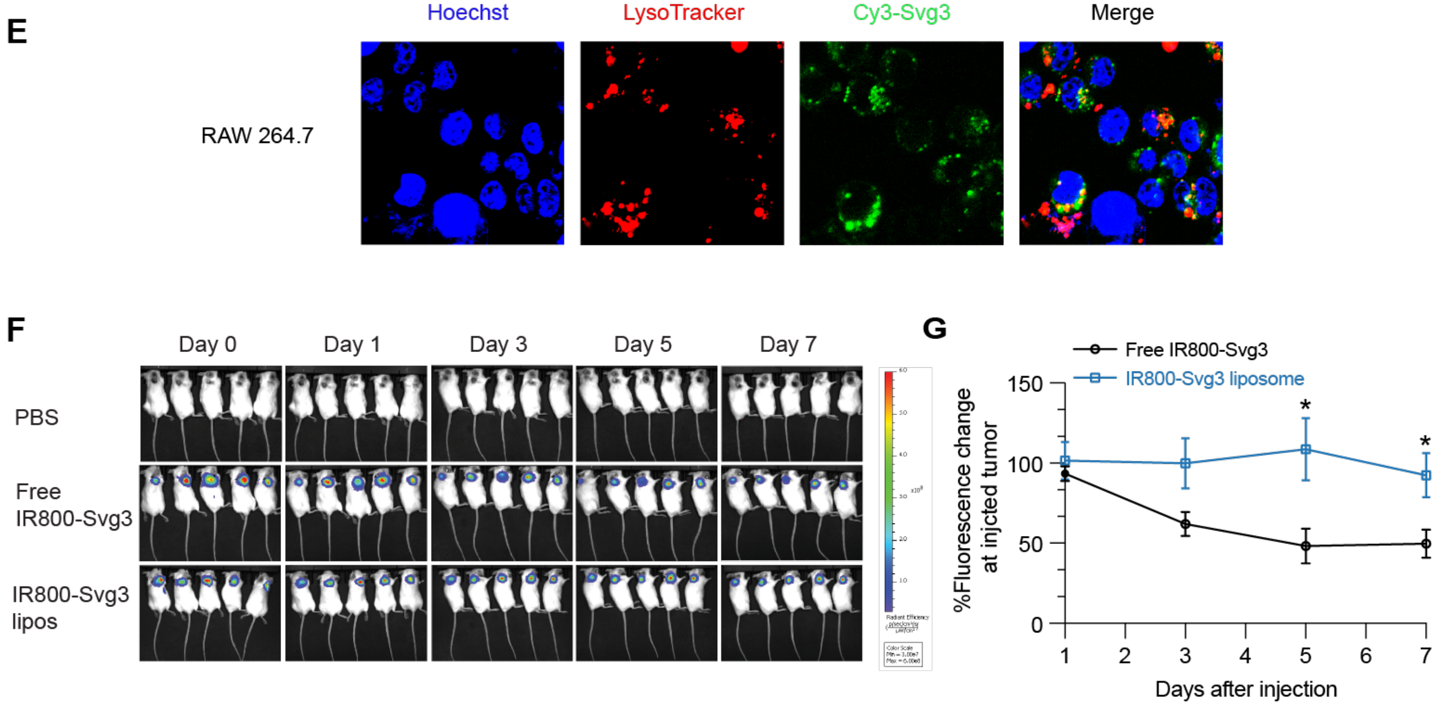

**Supplementary Figure 2. Liposomes readily loaded Svg3 for efficient delivery and retention of Svg3 in tissues and cells.** A) Characterization of the size, polydispersity index (PDI), zeta potential and encapsulation efficiency (EE%) of nanoparticles used in the study. B) Encapsulation efficiency of Svg3 liposomes evaluated by 2% agarose gel electrophoresis. (C, D) Median Cy3 fluorescence intensity from Cy3-Svg3 loaded in liposomes or LNPs. E) Confocal images of RAW 264.7 cells after incubated with liposomal Cy3-Svg3 for 5 h. F) IVIS imaging of 4T1 tumors after i.t. injection of liposomal IR800-Svg3. G) Fluorescence intensities of 4T1 tumors quantified from IVIS imaging results.

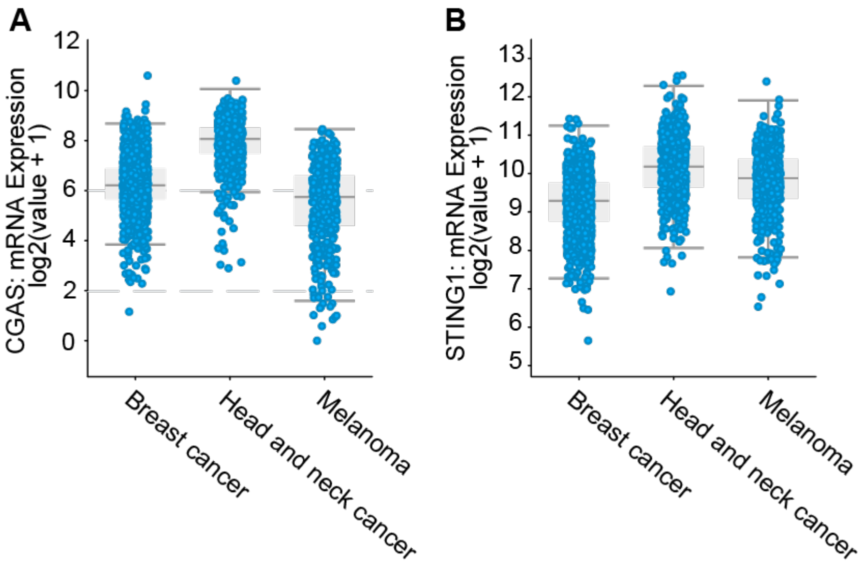

**Supplementary Figure 3.** The transcript levels of *cGAS* (A) and *STING* (B) in the indicated three types of human cancers. Human transcript data were downloaded from cBioPortal.

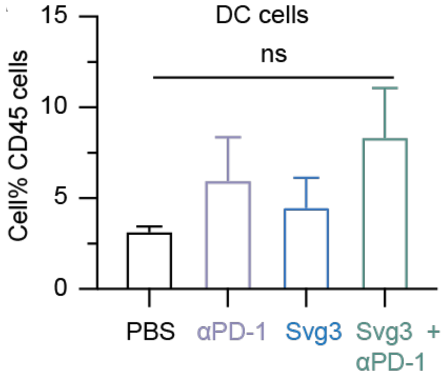

**Supplementary Figure 4.** Percentages of DC cells among CD45$^{+}$ cells in as-treated 4T1 tumors.

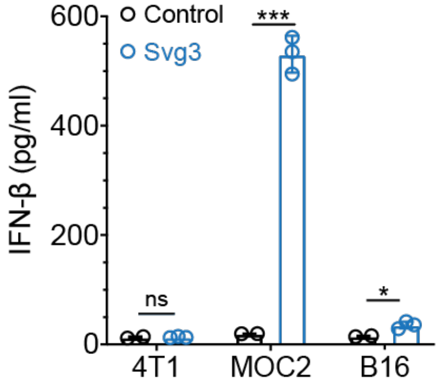

**Supplementary Figure 5. Svg3 elicited different levels of IFN-I responses in three types of murine tumor cells**, as shown by the IFN-β levels in the culture medium after Svg3 transfection (500 nM, 24 h).

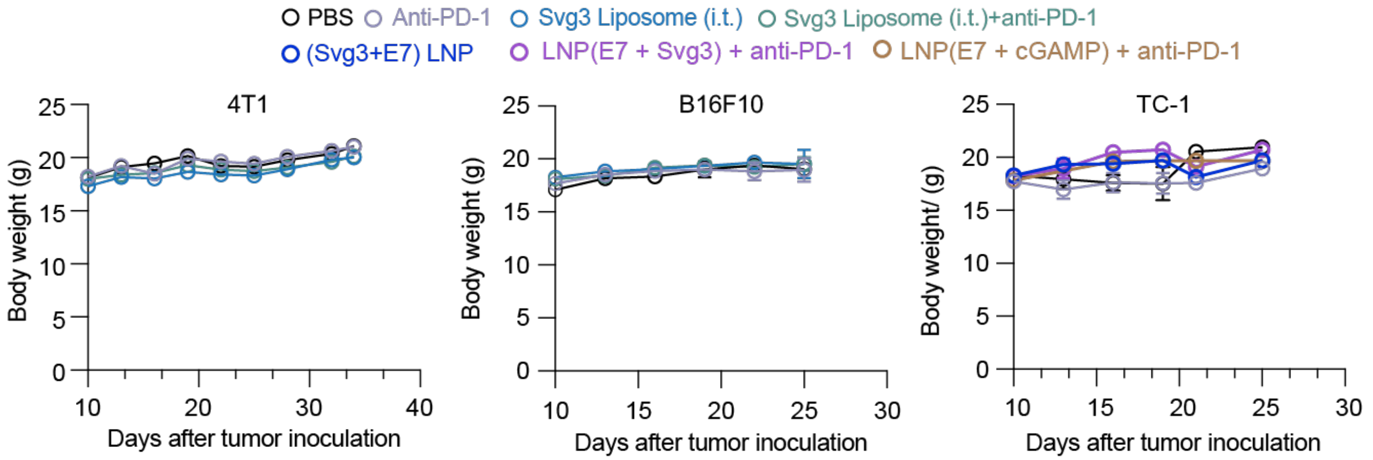

**Supplementary Figure 6.** Mouse body weights during the course of tumor therapy using the indicated treatments.

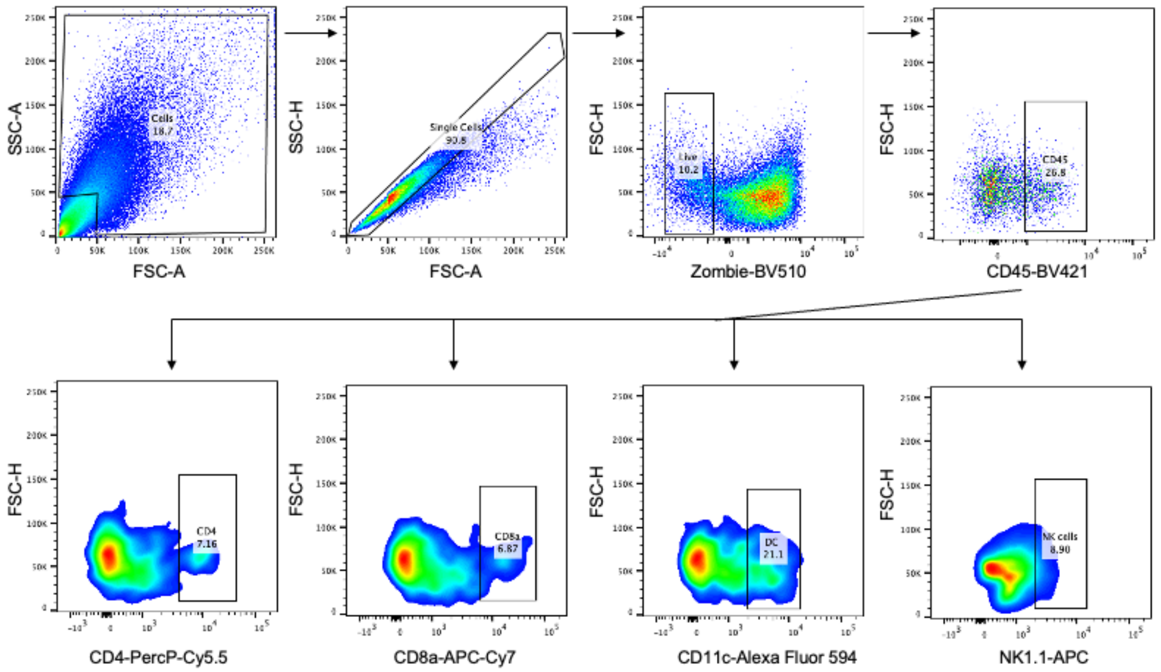

**Supplementary Figure 7.** Flow cytometry gating trees for the analysis of $CD4^+$ T cells, $CD8^+$ T cells, DCs, and NK cells in 4T1 tumor microenvironment.

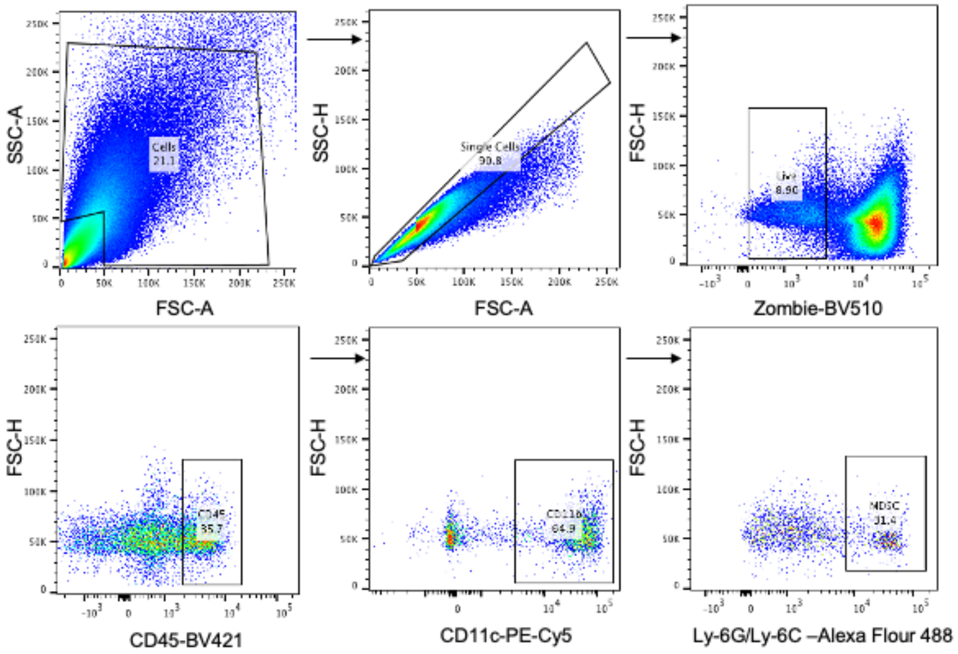

**Supplementary Figure 8.** Flow cytometry gating tree for the analysis of MDSCs in 4T1 tumor microenvironment analysis.

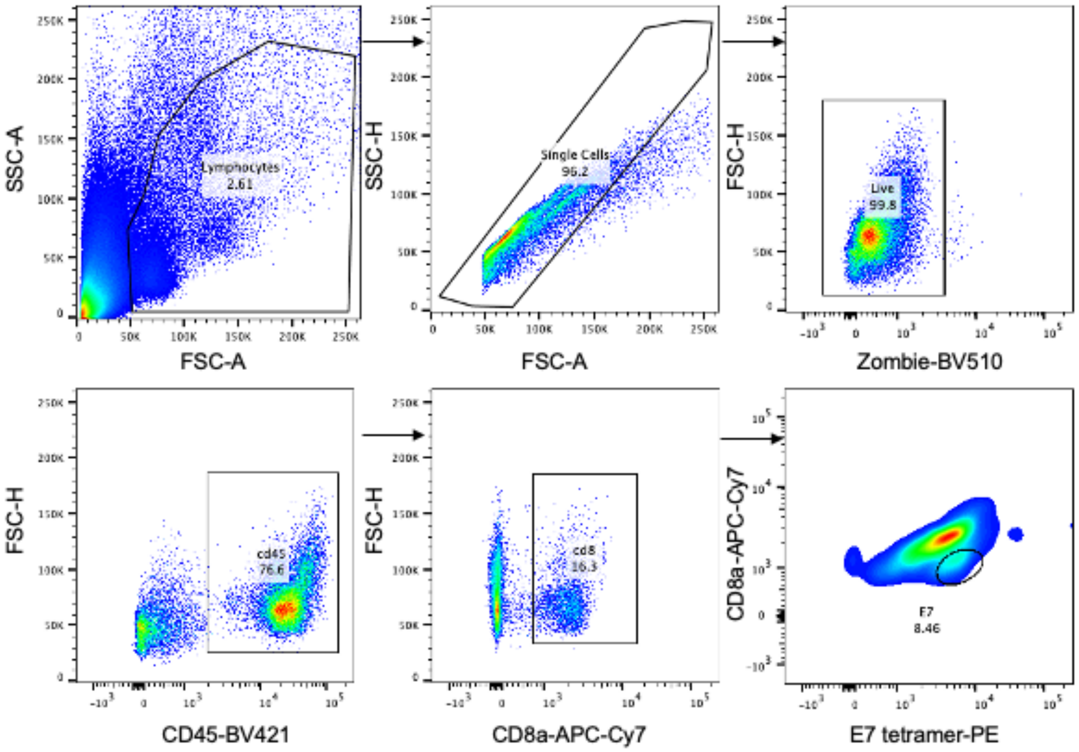

**Supplementary Figure 9.** Flow cytometry gating tree used for the analysis of E7 tetramer staining of PBMC $CD8^+$ T cells.

**Supplementary Table 1. DNA sequences used for oligonucleotide engineering and screening.**

| Name | Sequence (5'-3') |
|---|---|
| **Svg3** | CAG GGG GGA CCA CTC TTA AGC CTC AAG GGA AGC TGG GTT GAG GCT TAA GAG TGG TCC CGG GT |
| **Svg3-PS stem** | CAG GGG* G*G*A* C*C*A* C*T*C* T*T*A* A*G*C* C*T*C* A*AG GGA AGC TGG G*T*T* G*A*G* G*C*T* T*A*A* G*A*G* T*G*G* T*C*C* CGG GT |
| **Svg3-3dT** | CAG GGG GGA CCA CTC TTA AGC CTC AAG GGA AGC TGG GTT GAG GCT TAA GAG TGG TCC CGG GT /3InvdT/ |
| **Svg3-PS stem-3dT** | CAG GGG* G*G*A* C*C*A* C*T*C* T*T*A* A*G*C* C*T*C* A*AG GGA AGC TGG G*T*T* G*A*G* G*C*T* T*A*A* G*A*G* T*G*G* T*C*C* CGG GT/3InvdT/ |
| **Svc3** | CAC CCG GGA CCA CTC TTA AGC CTC AAC CCA AGC TCC GTT GAG GCT TAA GAG TGG TCC CCC CT |
| **Svg2** | CGGGGACCACTCTTAAGCCTCAAGCTGTTGAGGCTTAAGAGTGGTCCCGT |
| **Svg(3)** | GGGACCACTCTTAAGCCTCAAGGGAAGCTGGGTTGAGGCTTAAGAGTGGTCCC |
| **Svg4** | CAGGGGGACCACTCTTAAGCCTCAAGGAAGCTGGTTGAGGCTTAAGAGTGGTCCCGGT |
| **ISD (InvivoGen)** | 5'-TACAGATCTACTAGTGATCTATGACTGATCTGTACATGATCTACA-3' |
| | 3'-ATGTCTAGATGATCACTAGATACTGACTAGACATGTACTAGATGT-5' |
| **DNA_10 bp** | TAC TCC AAG CAA GCC CGC TTG GAG TAG TCT TG |
| **DNA_14 bp** | GTT CTG TTG AGG ATT TAC GGA CAC AAC CGT AAA TCC TCA A |
| **DNA_18 bp** | GTT CTG TTG AGG ATT TAC GAG GTCAC ACA AGACC TCG TAA ATC CTC AA |
| **DNA_20 bp** | GTT CTG TTG AGG ATT TAC GAG GTCGTAC ACA AACGACC TCG TAA ATC CTC AA |
| **DNA_21 bp** | CAC CCC GGA CCA CTC TTA AGC CTC AAC CCA AGC TCC GTT GAG GCT TAA GAG TGG TCC CCC CT |
| **DNA_22 bp** | GTT CTG CTTTG AGG ATT TAC GAG GTCGTAC ACA AACGACC TCG TAA ATC CTC AAAG |
| **DNA_24 bp** | GTT CTG AGCTTTG AGG ATT TAC GAG GTCGTAC ACA AACGACC TCG TAA ATC CTC AAAGCT |

**Supplementary Table 2. Primer sequences used for qPCR.**

| ***Human genes*** | 5'-3' |
|---|---|
| **Gapdh-Forward** | GGAGCGAGATCCCTCCAAAAT |
| **Gapdh-Reverse** | GGCTGTTGTCATACTTCTCATGG |
| **Ifna2-Forward** | GCTTGGGATGAGACCCTCCTA |
| **Ifna2-Reverse** | CCCACCCCCTGTATCACAC |
| **Ifnb-Forward** | ATGACCAACAAGTGTCTCCTCC |
| **Ifnb-Reverse** | GGAATCCAAGCAAGTTGTAGCTC |
| **Il6-Forward** | ACTCACCTCTTCAGAACGAATTG |
| **Il-Reverse** | CCATCTTTGGAAGGTTCAGGTTG |
| **Cxcl10-Forward** | GTGGCATTCAAGGAGTACCTC |
| **Cxcl10-Reverse** | TGATGGCCTTCGATTCTGGATT |
| ***Mouse genes*** | |
| **Gapdh-Forward** | CTTTGTCAAGCTCATTTCCTGG |
| **Gapdh-Reverse** | TCTTGCTCAGTGTCCTTGC |
| **Tnfa-Forward** | GGTGCCTATGTCTCAGCCTCTT |
| **Tnfa-Reverse** | GCCATAGAACTGATGAGAGGGAG |
| **Il6-Forward** | GAGGATACCACTCCCAACAGACC |
| **Il-Reverse** | AAGTGCATCATCGTTGTTCATACA |
| **Cxcl10-Forward** | ATCATCCCTGCGAGCCTATCCT |
| **Cxcl10-Reverse** | GACCTTTTTTGGCTAAACGCTTTC |

**Supplementary Table 3. Peptide sequences.**

| | |
|---|---|
| **SIINFEKL** | SIINFEKL |
| **HPV-16 $E7_{43-62}$** | GQAEPDRAHYNIVTFCCKCD |

**Supplementary Table 4. Details of antibodies used in this study.**

| Targets | Fluorochrome | Clone | Vendor | Catalogue # |
|---|---|---|---|---|
| **CD45** | Brilliant Violet 421 | 30-F11 | BioLegend | 103133 |
| **CD11c** | Alexa Fluor 594 | N418 | BioLegend | 117346 |
| **CD11b** | FITC | M1/70 | BioLegend | 101205 |
| **CD8a** | APC/Cy7 | 53-6.7 | BioLegend | 100713 |
| **CD4** | PerCP/Cy5.5 | GK1.5 | BioLegend | 100433 |
| **CD103** | Brilliant Violet 605 | 2.00E+07 | BioLegend | 121433 |
| **CD205** | PE/Cy7 | NLDC-145 | BioLegend | 138209 |
| **F4/80** | APC/Cy7 | BM8 | BioLegend | 123117 |
| **CD11b** | PE/Cy5 | M1/70 | BioLegend | 101209 |
| **NK1.1** | APC | S17016D | BioLegend | 156505 |
| **CD3** | PerCP/Cy5.5 | 17A2 | BioLegend | 100217 |
| **CD44** | PE/Cy5 | IM7 | BioLegend | 103009 |
| **CD62L** | FITC | MEL-14 | BioLegend | 104405 |
| **CD25** | FITC | 3C7 | BioLegend | 101907 |
| **FoxP3** | Alexa Fluor 647 | MF-14 | BioLegend | 126407 |
| **CD279 (PD-1)** | N/A | RMP1-14-CP162 | Bio X Cell | CP162 |
